# Oligogenic combinations of rare variants influence specific phenotypes in complex disorders

**DOI:** 10.1101/2021.10.01.462832

**Authors:** Vijay Kumar Pounraja, Santhosh Girirajan

**Affiliations:** Department of Biochemistry and Molecular Biology, Pennsylvania State University, University Park, PA 16802; Bioinformatics and Genomics Graduate Program, The Huck Institute of the Life Sciences, University Park, PA 16802; Department of Anthropology, Pennsylvania State University, University Park, PA 16802

**Keywords:** oligogenic, complex disorders, autism, rare variants

## Abstract

Genetic studies of complex disorders such as autism and intellectual disability (ID) are often based on enrichment of individual rare variants or their aggregate burden in affected individuals compared to controls. However, these studies overlook the influence of combinations of rare variants that may not be deleterious on their own due to statistical challenges resulting from rarity and combinatorial explosion when enumerating variant combinations, limiting our ability to study oligogenic basis for these disorders. We present a framework that combines the apriori algorithm and statistical inference to identify specific combinations of mutated genes associated with complex phenotypes. Our approach overcomes computational barriers and exhaustively evaluates variant combinations to identify non-additive relationships between simultaneously mutated genes. Using this approach, we analyzed 6,189 individuals with autism and identified 718 combinations significantly associated with ID, and carriers of these combinations showed lower IQ than expected in an independent cohort of 1,878 individuals. These combinations were enriched for nervous system genes such as *NIN* and *NGF*, showed complex inheritance patterns, and were depleted in unaffected siblings. We found that an affected individual can carry many oligogenic combinations, each contributing to the same phenotype or distinct phenotypes at varying effect sizes. We also used this framework to identify combinations associated with multiple comorbid phenotypes, including mutations of *COL28A1* and *MFSD2B* for ID and schizophrenia and *ABCA4, DNAH10* and *MC1R* for ID and anxiety/depression. Our framework identifies a key component of missing heritability and provides a novel paradigm to untangle the genetic architecture of complex disorders.

**SIGNIFICANCE:** While rare mutations in single genes or their collective burden partially explain the genetic basis for complex disorders, the role of specific combinations of rare variants is not completely understood. This is because combinations of rare variants are rarer and evaluating all possible combinations would result in a combinatorial explosion, creating difficulties for statistical and computational analysis. We developed a data mining approach that overcomes these limitations to precisely quantify the influence of combinations of two or more mutated genes on a specific clinical feature or multiple co-occurring features. Our framework provides a new paradigm for dissecting the genetic causes of complex disorders and provides an impetus for its utility in clinical diagnosis.

## INTRODUCTION

Recent human population growth has led to a rapid increase in the load of rare variants affecting functionally important regions of the genome^1–3^. Thus, rare variants are collectively more abundant in the population compared to common variants, many of which confer significant risk for neurodevelopmental disorders such as autism and intellectual disability^4^. In fact, recent studies have directly implicated rare damaging mutations that are very recent or *de novo* in more than one hundred genes towards neurodevelopmental disorders^5–7^. The ability to establish robust associations between rare variants of high effect size and complex disease has made this class of variants the primary focus of recent studies. However, a much larger class of rare and variably expressive variants that are individually less deleterious but, in combination, exert large effects towards disease is often overlooked. Variants in this category are often transmitted across generations without adverse effects on their carriers until they encounter other similar variants that, when combined, lead to genetic interactions conferring a higher risk for disease than their individual risks^8,9^. While this phenomenon underpins oligogenic models proposed over the years, studies so far have not focused on detecting combinatorial effects of specific sets of rare variants towards disease phenotypes^10–13^.

Identifying the effects of specific combinations of rare variants towards disease etiology has been challenging for many reasons. *First*, combinations of rare variants are rarer, and extremely large cohorts are required to observe even a few recurrent instances of specific variant combinations^14^. Prior studies of oligogenic models for rare variants evaded this problem by aggregating variant information at the sample level and comparing the overall burden of rare variants between groups of individuals (such as cases and controls)^6,7,15,16^. *Second*, the combinatorial explosion resulting from even a small set of rare variants makes it difficult to exhaustively evaluate all combinations. While sophisticated frameworks such as network analysis and machine learning provide powerful tools to model the composite effects of thousands of variables on a complex system and predict emergent behaviors and quantitative outcomes, adapting them to exhaustively search and delineate the effects of specific combinations of variables is daunting^17,18^. Furthermore, incorporating an efficient search tool into these frameworks and extending them to detect higher-order combinatorial effects would be nearly impossible. *Third*, even when all combinations of rare variants could be exhaustively evaluated within a large cohort, there is a lack of methods that are sensitive enough to detect small differences between comparison groups to establish statistical significance. Therefore, an alternate approach that is highly flexible, scalable, and sensitive is necessary to address computational and statistical challenges associated with assessing rare variant combinations.

Here, we present a combinatorial framework called *RareComb* that couples the apriori algorithm^19^ with binomial tests to overcome the limitations of data sparsity and high dimensionality, and systematically analyzes patterns of rare variants between groups of interest to identify specific combinations that are significantly associated with phenotypes^20^. We apply our analysis framework to a discovery cohort of 6,189 children with autism to identify genetic interactions involving pairs and triplets of mutated genes that are enriched in individuals with intellectual disability compared to individuals without intellectual disability. We demonstrate that the carriers of mutations in these specific gene pairs and triplets within an independent cohort of 1,878 children have significantly lower-than-expected intelligence quotient (IQ) scores. We also demonstrate the adaptability of our framework by leveraging it to identify mutated gene pairs and triplets significantly associated with two or more comorbid phenotypes among children with autism. Finally, we show how this generalizable and modular framework can be easily extended to identify higher order interactions beyond pairs and triplets of variants. Our stand-alone framework does not depend on *a priori* knowledge and can detect rare patterns from high-dimensional genetic data to generate interpretable results, making it readily applicable for analyzing cohorts of all size ranges to dissect the genetic basis of complex disorders.

## RESULTS

We hypothesized that two or more genes disrupted simultaneously by rare deleterious mutations contribute to a highly penetrant phenotype, as in an oligogenic model, or lead to a more severe phenotype than when each of the same genes are disrupted individually. We developed *RareComb* as a framework that combines data mining and statistical analysis to identify specific combinations (such as pairs, triplets, etc.) of rare variants that show significant associations with one or more phenotypes. *RareComb* analyzes an ‘*n*×*p*’ sparse Boolean matrix with ‘*p*’ genes in ‘*n*’ individuals in three discrete steps (**Figure 1**). *First*, it applies the apriori algorithm independently in cases and controls to enumerate the frequency of all simultaneously mutated combinations that meet a pre-set minimum frequency threshold (**Supp. Figure 1**). *Second*, for each qualifying combination of variants, the method derives the expected frequency of simultaneously observing mutations in the constituent genes under the assumption of independence. It then independently quantifies the magnitude of deviation of the observed from the expected frequencies using binomial tests in cases and controls, and uses multiple-testing adjusted p-values to identify combinations that are statistically enriched in cases but not in controls. *Finally*, the method calculates effect sizes using Cohen’s d and statistical power at 1% and 5% significance thresholds, to enable prioritization of a high confidence set of combinations that contribute to the phenotype in an oligogenic manner.

**Figure 1:**
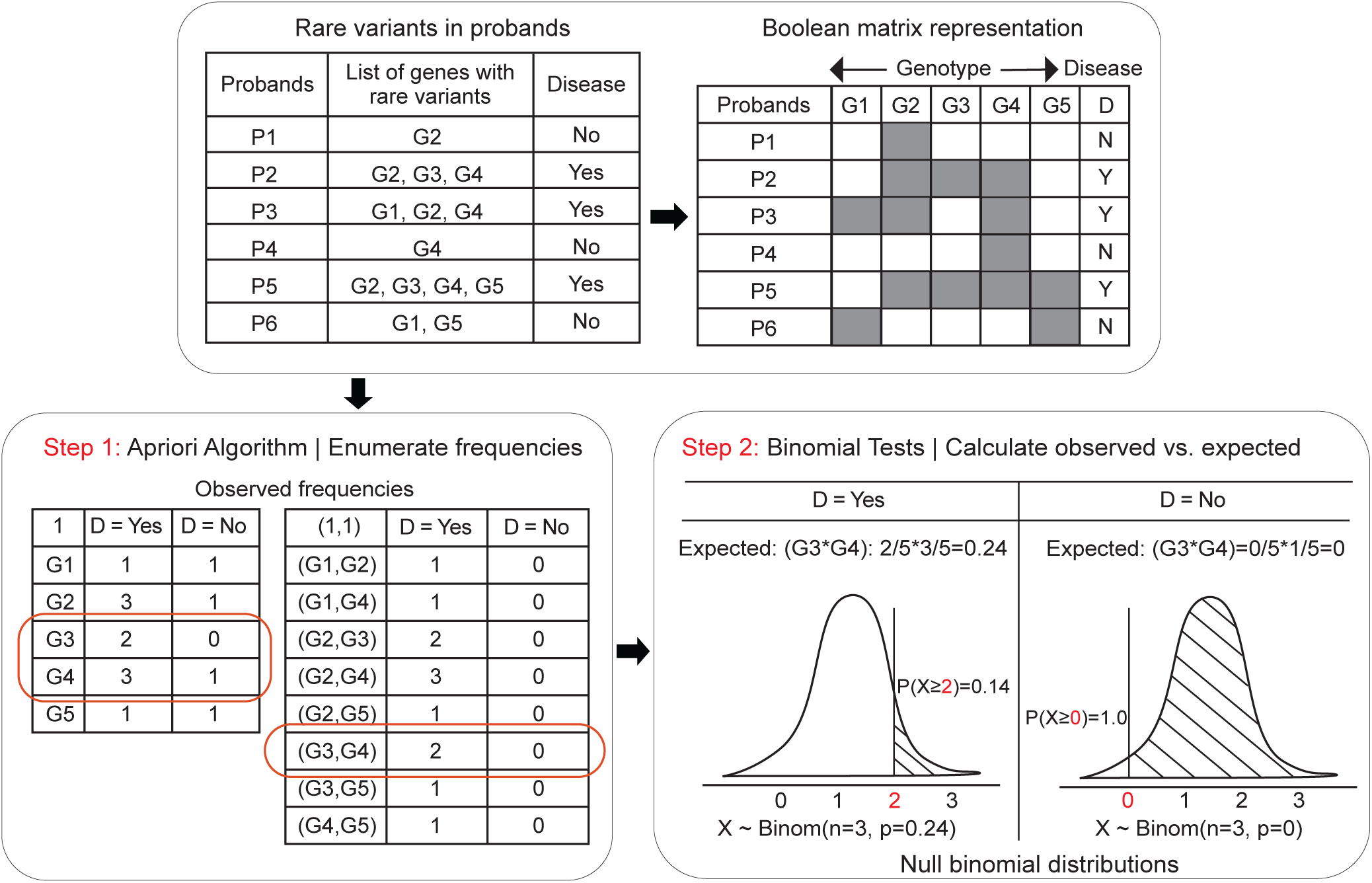
Conceptual overview of combinatorial analyses using *RareComb*. A Boolean representation of genotype (mutated genes, G1, G2, etc) and disease status for probands (P1, P2, etc) is shown. In *step 1*, the apriori algorithm is applied to the Boolean input matrix to calculate the frequencies of individual (for example, G1) and simultaneous occurrences of events (G1 and G2) that meet the user-specified criteria, including the size of combinations (pairs, triplets, etc.) and minimum frequency threshold of simultaneous occurrences. In *step 2*, independently in case and control groups, for each combination, the binomial test is applied to compare the observed frequency of simultaneous occurrence of events with its corresponding null binomial distribution of the expected frequencies calculated under the assumption of independence. Binomial test for gene pair G3 and G4 is shown as an example.

### *RareComb* identifies oligogenic combinations associated with ID and autism

We sought to identify pairs and triplets of mutated genes that are significantly associated with intellectual disability (ID) phenotypes by analyzing 6,189 affected individuals from the SPARK^21^ cohort for discovery and 1,878 affected individuals from the SSC^22^ cohort for validation. To facilitate cross-cohort comparison, we identified 10,217 rare variants (MAF≤1%) that were predicted to be deleterious by multiple methods and observed in both cohorts, and aggregated these variants to genes for the analysis (see **Methods**). We first categorized 1,215 probands from the SPARK cohort diagnosed with ID as cases and 4,974 probands without ID as controls (**Figure 2A**). We then applied *RareComb* to cases after constraining it to only evaluate those gene combinations in which simultaneous mutations are observed in at least five probands. We identified 25,602 pairs involving 1,956 mutated genes in cases that were observed at a higher frequency than expected under the assumption of independence. Similarly, analyzing the controls using only the 1,956 genes mutated in cases, *RareComb* identified 148 pairs of mutated genes that were significantly enriched in cases but not in controls (**Supp. Table 1)**, with moderate to high effect sizes (Cohen’s d, 0.08-0.15) and adequate statistical power (70%-100% at 5% significance threshold) (**Supp. Figure 2**). These 148 gene pairs belonged to 142 probands, with 74% (105/142) of them carrying more than one significant pair. These observations suggest that an individual can carry multiple combinations, each contributing to the same phenotype at varying effect sizes (**Supp. Figure 3**).

**Figure 2:**
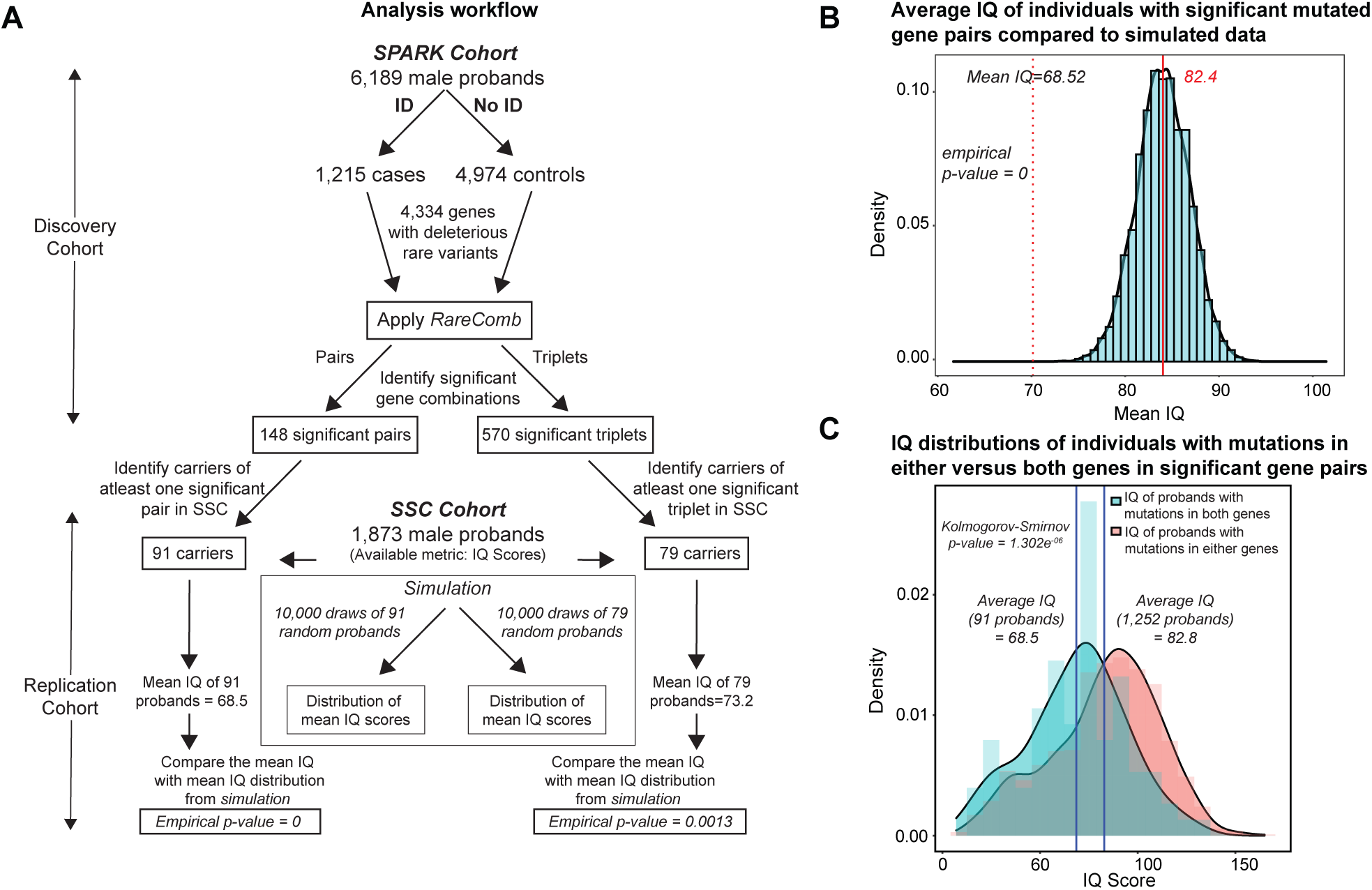
Combinations of rare variants contributing to intellectual disability (ID) phenotype. **(A)** An outline of the approach used to identify and validate mutated gene pairs and triplets enriched in probands with ID is shown. We tested whether mutated gene pairs identified as significant in one cohort (SPARK) are also associated with severe phenotypes in an independent cohort (SSC). To test this, we obtained the mean IQ score of individuals from the SSC cohort carrying significant combinations identified from the SPARK cohort. Empirical p-values were then calculated based on the deviation of the mean IQ from the distribution of mean IQ scores obtained from 10,000 random draws in the simulation. **(B)** The mean IQ of individuals with mutated gene pairs in the SSC cohort was significantly lower (empirical p-value=0) when compared to the distribution of mean IQ scores obtained from the simulation. **(C)** Histogram shows the distributions of IQ scores of SSC probands who carried mutations in either genes versus both constituent genes of the significant gene pairs. The distributions were significantly different from each other (p-value = 1.302×10^−6^, Kolmogorov-Smirnov test).

We next sought to validate the association of these 148 mutated gene pairs towards intellectual disability. We hypothesized that if the association of the gene pairs with ID in the SPARK cohort were truly significant, carriers of mutations in those gene pairs would tend to have lower than average IQ scores in the independent SSC cohort. We found that 90 of the 148 significant pairs identified in the SPARK cohort were observed in at least one proband in the SSC cohort. These 90 mutated gene pairs were carried by 91 unique probands, whose average full-scale IQ scores (average IQ=68.52) were lower than those of all ascertained probands in the SSC cohort (average IQ=86). To assess the significance of this result, we performed 10,000 random draws of 91 probands from the SSC cohort to generate a simulated distribution of their average IQ scores. The average IQ of carriers of mutated gene pairs (average IQ=68.52) was significantly lower than the overall distribution of average IQ derived from simulations (average IQ ranged from 73 to 92; empirical p=0) (**Figure 2B**). Furthermore, the average IQ of the 91 SSC probands with both mutated genes was significantly lower than the average IQ of 1,252 carriers of mutations in only one of the two genes (68.5 versus 82.8; Kolmogorov-Smirnov p = 1.302×10^−16^) (**Figure 2C**). When each of the 90 combinations was evaluated individually, carriers of mutations in both genes for 73% (66/90) of the combinations showed lower IQ than individuals with mutations in individual genes of the same combination, with 39/90 remaining significant after multiple testing correction (**Supp. Table 2; Supp. Figure 4**). These results provide evidence for synergistic effects of deleterious mutations within specific pairs of genes towards ID phenotypes.

We also applied *RareComb* to identify gene triplets associated with intellectual disability using the two cohorts and repeated the simulations to identify 1,593 significant combinations in the SPARK cohort. We selected 570 high-confidence triplets (with ≥90% statistical power at 5% significance threshold; **Supp. Table 3**) and found that 79 probands in the SSC cohort carried at least one of these deleterious triplets. The average IQ score of individuals carrying significant gene triplets (average IQ score=73) was significantly lower than a distribution of average IQ scores from 10,000 draws of 79 SSC probands (average IQ score=82.5; min=72, max=94; empirical p=0.0011; see **Supp. Figure 5**). This result reiterated that carriers of mutations in the significant gene combinations have lower IQ than a random group of probands. Our results also demonstrate the ability of the framework to identify higher order combinations of mutations that are significantly associated with specific phenotypes in individuals with complex disorders.

### Oligogenic combinations are enriched for specific inheritance patterns

As individual variants can arise *de novo* or be inherited maternally or paternally, variants in pairs of genes can have six possible patterns of transmission (**Supp. Figure 6A**). We identified a total of 926 occurrences of the 148 pairs of mutated genes enriched among SPARK probands with ID (n=142 probands), of which inheritance could be determined without ambiguity for 887 instances. We found that one variant occurred *de novo* and the other variant was inherited from the mother in 244/887 instances (27.5%). Similarly, both mutated genes were inherited from the mother in 226/887 instances (25.4%) or occurred *de novo* in 221/887 instances (24.9%), while the remaining fraction (∼22%) of variant pairs were either inherited from both parents, inherited from the father, or transmitted *de novo* and paternally. To assess the significance of our observations, we performed simulations to establish a baseline expectation of proportions for each category of parental inheritance pattern. We selected 926 pairs of genes in 1000 random draws of all possible mutated gene pairs among SPARK probands and calculated the fraction of instances that fell into each of the six transmission categories. The observed proportion was higher than the simulated proportions for instances when both variants occurred *de novo* (24.9% versus 17%, empirical p=0) and when one variant was *de novo* and the other was inherited maternally (27.5% versus 25%, p=0.028) (**Figure 3A**). We repeated this analysis for 7,596 children affected with autism in the SPARK cohort compared to 11,740 unaffected parents and identified 110 gene pairs significantly associated with autism (**Supp. Table 4**). Similar to the results obtained for the ID phenotype, we found that both variants of a gene pair were more likely to occur *de novo* (24% versus 18%, empirical p=0) or one variant occurring *de novo* and the other inherited maternally (33% versus 26%, p=0) than expected based on simulation studies (**Supp. Figure 7**). The enrichment of *de novo* or maternally inherited variants for significant gene pairs aligns with published reports that severely affected children tend to carry multiple *de novo* mutations or inherit pathogenic rare variants from mildly affected or unaffected carrier mothers^16,23,24^.

**Figure 3:**
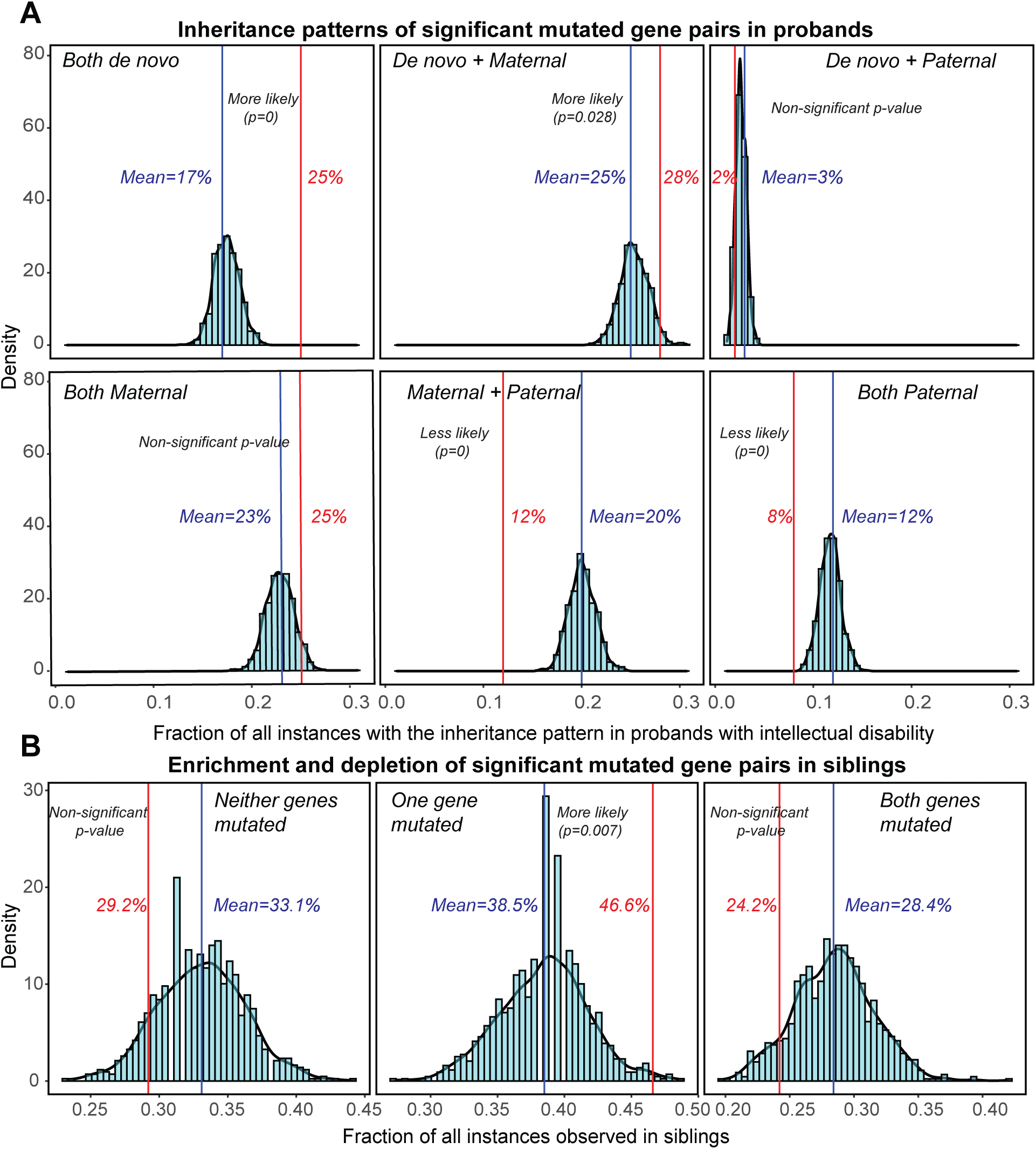
Analysis of parental and sibling inheritance patterns of significant gene pairs associated with ID. **(A)** Fraction of all instances of significant gene pairs observed within each of the six possible parental inheritance patterns (red) compared against 1,000 simulations is shown (blue). During each simulation, random mutated gene pairs from the SSC cohort were selected, the inheritance status of the mutations was identified, and the fraction of those instances belonging to one of the six pre-defined categories was calculated. Comparing the observed fractions with the simulated fractions indicate statistical enrichment for two specific inheritance patterns based on empirical p-values: both variants being *de novo*, and one variant being *de novo* and the other transmitted from the mother. **(B)** Histograms show the carrier status of significant gene pairs in siblings of carrier probands (red) compared against 1,000 simulations (blue). Among significant pairs, both genes were mutated in only 24.2% of all siblings (compared to 28.4% in simulations), whereas one of the two genes was mutated in 46.6% of all siblings (compared to 38.5% in simulations). These results show that mutations are more likely to be observed in just one of the two genes within the gene pairs and are less likely to be observed simultaneously in siblings of carrier probands.

We then assessed whether the mutated gene pairs associated with ID were also found in siblings of carrier probands. Restricting our analysis to families with unaffected siblings whose probands had mutations in ID-enriched gene pairs, we found that both variants were present in the corresponding sibling for only 53/219 (24.2%) instances of gene pairs, while 102/219 (46.6%) had variants in only one of the two genes and 64/219 (29.2%) instances had no variants in either of the genes in the siblings (**Supp. Figure 6B)**. Using simulations, we found a significantly higher proportion of instances with only one of the two variants present in siblings compared to the expected values (46.6% versus 38.5%, p=0.007). Furthermore, the proportion of observed instances with neither of the variants present in siblings (29.2% versus 33.1%, empirical p=0.098) or both variants present in siblings (24.2% versus 28.4%, p=0.079) was lower than expected (**Figure 3B)**. The observation that only a small fraction of unaffected siblings carried both mutated gene pairs suggests a strong association of these gene pairs with ID phenotypes. These results suggest that mutations in pairs of genes significantly associated with a severe phenotype in probands are more likely to occur individually than simultaneously in unaffected siblings of the same family.

### Genes forming oligogenic combinations are distinct from canonical autism genes

We expanded our analysis to include all 16,556 mutated genes in the SPARK cohort, as opposed to genes with mutations present in both the SPARK and SSC cohorts, and identified 52 significant gene pairs (**Supp. Table 5)** and 230 triplets associated with the ID phenotype (with ≥90% statistical power at 1% significance threshold; **Supp. Table 6)**. Due to the expanded search space, the mutated gene pairs showed more significant p-values from the binomial tests when compared to those obtained from the more restricted set of variants overlapping both SPARK and SSC cohorts (**Supp. Figure 8**). Mutated genes within these combinations included several genes related to nervous system development, such as *NIN, HDC, NGF*, and *BRD8*. Furthermore, 5/52 pairs and 59/230 triplets contained at least one gene associated with autism in the SFARI database, including *FGFR1*, associated with multiple disorders including Kallmann syndrome^25^ and Pfeiffer syndrome^26^; *RELN*, associated with temporal lobe epilepsy^27^; *SYNE1*, associated with spinocerebellar ataxia^28,29^; and *PNPLA7*, associated with autism and ID^30^. Thus, most genes forming combinations are not involved in canonical autism or ID disorders, suggesting synergistic effects of these genes without prior association to disease.

We also performed gene ontology enrichment analysis for genes within the combinations and identified seven out of nine significantly enriched GO terms to be exclusively associated with nervous system-related functions, including synthesis and metabolism of catecholamines, axon/neuron regeneration, and neuron generation and differentiation (**Supp. Figure 9**)^31^. Furthermore, the differences in the type and specificity of GO terms enriched for significant pairs versus triplets were apparent, with genes forming pairs involved in nervous system function and genes forming triplets associated with both nervous system as well as other biological processes. We next assessed the enrichment and depletion of Human Phenotype Ontology (HPO) terms for genes forming significant pairs towards ID phenotypes^32^. *First*, we calculated the fraction of all 4,484 genes within the HPO database associated with each HPO term. *For example*, 30% (1,366/4,484) of all genes in HPO were associated with ID. We compared these expected values calculated for each HPO term with the corresponding fractions observed within the 95 genes forming 52 ID-associated pairs using binomial tests. Interestingly, genes associated with HPO terms related to neurodevelopmental phenotypes, such as ID, global developmental delay, seizure, and microcephaly, were significantly depleted within the set of 95 genes forming gene pairs (**Supp. Table 7**). *Next*, we evaluated whether genes within each of the 52 significant pairs shared one or more common HPO phenotype or disease. Of the 52 pairs, only one pair (*DNASE1 & MTR*) shared an HPO phenotype (“epilepsy”). This was significantly lower than the expected value obtained from the distribution of the number of shared HPO phenotypes between all possible pairs of genes in the HPO database (1/52, 1.9% ID gene pairs compared to 31.5% of all HPO gene pairs shared one HPO phenotype, p= 2.2×10^−16^; one-sided binomial test) (**Supp. Figure 10; Supp. Table 8**). We note that the 4,484 genes within HPO are potentially biased towards well-studied disorders, making pairs of genes drawn from HPO more likely to share phenotypes than random pairs of genes from the genome. Overall, GO and HPO analyses show that genes forming oligogenic combinations are involved in neuronal processes but have not been previously connected to neurodevelopmental phenotypes, indicating the novelty of the associations between these genes and ID phenotypes.

### Identifying variant combinations towards specific patterns of comorbid phenotypes

We adapted our framework to identify significant associations of two or more genotypes with multiple comorbid phenotypes. To identify novel comorbid associations, we eliminated phenotypes that were highly correlated with each other, such as ADHD and reading disorder^33^. We analyzed variant profiles of 6,189 autism probands from the SPARK cohort with records of comorbid features, including 1,215 individuals with ID, 1,825 with anxiety and depression, and 332 with schizophrenia features. We assessed for significant co-occurrences of two or more mutated genes with two or more of the above phenotypes (**Figure 4**). Using one-tailed binomial tests to compare the observed frequency of combinations of genotypes and phenotypes to the expected frequency, we first identified 169 significant associations between pairs of mutated genes and two comorbid phenotypes as well as 82 combinations of three mutated genes and two comorbid phenotypes (**Supp. Tables 9 & 10**). As some of these significant genotype-phenotype combinations can be confounded by high degree of co-occurrence of mutated genes, we next calculated genotype-only p-values using binomial tests for all significant genotype-phenotype associations. For 32/169 combinations of two mutated genes and two comorbid phenotypes and 5/82 combinations of three mutated genes and two comorbid phenotypes, the composite genotype-phenotype p-values were significant while genotype-only p-values were not significant, suggesting stronger associations between these variant combinations and phenotypes. *For example*, even when variants in genes *COL28A1* and *MFSD2B* did not co-occur more frequently than expected under the assumption of independence, these mutated genes co-occurred more frequently than expected among probands with ID and schizophrenia phenotypes. Loss-of-function and rare missense mutations in *COL28A1* have been reported in individuals with autism^34,35^, and *MFSD2A, a* paralog of *MFSD2B*, has been directly implicated in an autosomal recessive disorder associated with progressive microcephaly, spasticity and brain imaging abnormalities^36^. Similarly, we found *ARVCF* and *FAT1* to be significantly associated with ID and schizophrenia, with *ARVCF* mapping within the 22q11.2 DiGeorge syndrome region^37^, while rare *de novo* mutations in *FAT1* being associated with autism and schizophrenia^6,38^. Finally, we found that the mutations in genes *ABCA4, DNAH10* and *MC1R* significantly co-occurred in individuals with ID and anxiety/depression phenotypes. These results demonstrate the utility of identifying higher-order associations between genotypes and phenotypes in complex disorders such as autism.

**Figure 4:**
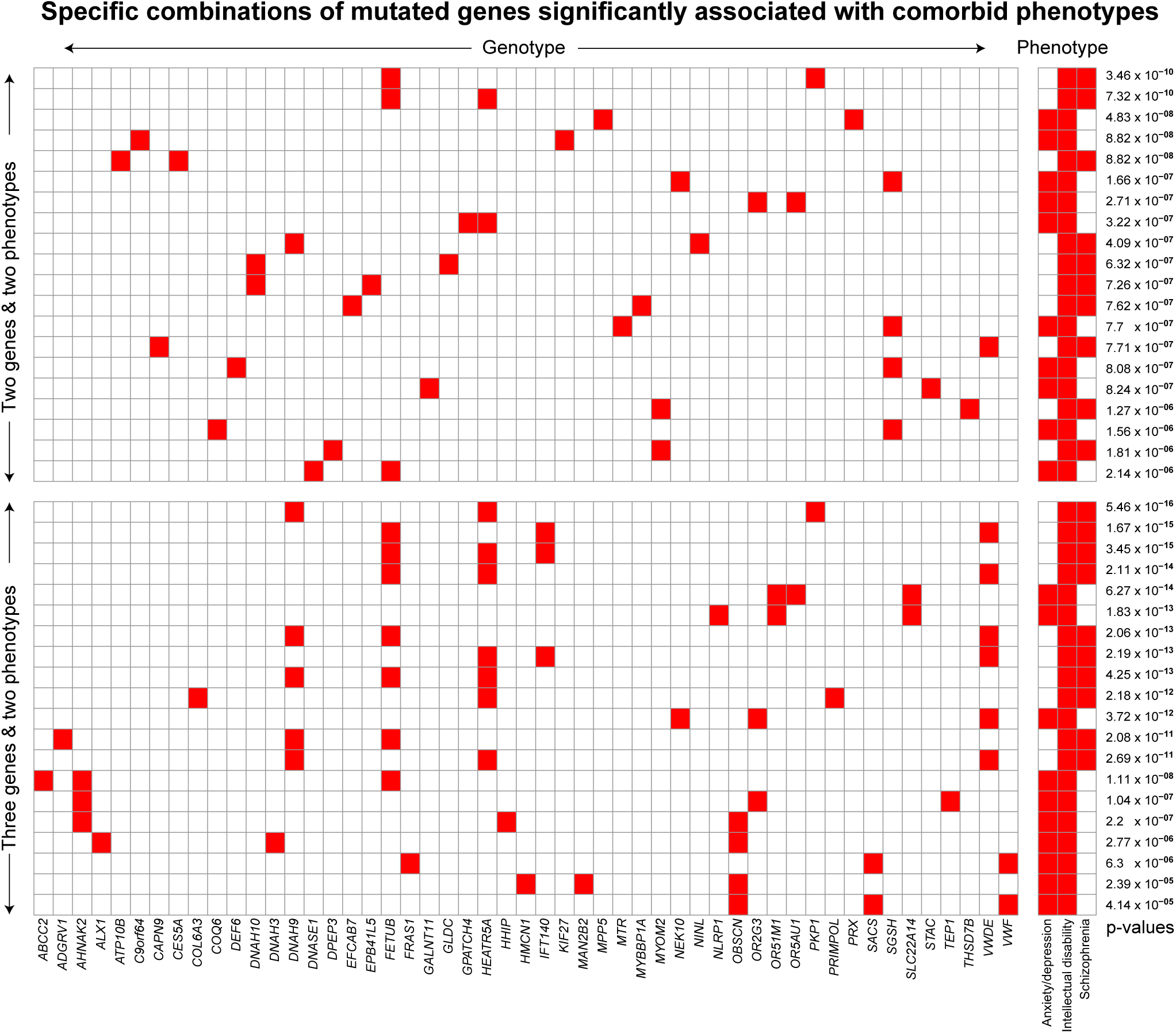
Analysis of comorbid phenotypes using *RareComb*. We analyzed the genotypes of probands with anxiety/depression, ID, or schizophrenia. The heatmap shows combinations of two or three mutated genes that were significantly enriched in individuals with specific patterns of comorbid phenotypes compared to the expected frequency under the assumption of independence.

## DISCUSSION

Current rare variant analysis strategies are geared towards either searching for individual variants of high effect size whose influence on the phenotype is evident, such as *de-novo* gene-disruptive mutations, or comparing rare variant burden to explain collective effects on phenotypes^7,39,40^. The wider space between these two extremes of the analysis spectrum that involves combinations of rare variants has largely remained understudied. Although digenic diseases and multi-hit models of complex diseases have been used to provide post-hoc explanations for an observed phenomenon, they are not equipped to serve as a framework to actively search and identify rare variant combinations that fit oligogenic models for specific phenotypes^9,12,13^. While machine learning has become the de-facto approach for disease outcome predictions, the lack of holy-grail predictors and reduced interpretability due to data sparsity makes it less fit to detect combinatorial effects^17^. In addition, the common practice of evaluating feature importance metrics of machine learning classifiers falls short of the objective to identify combinations of features that exert higher effect on the phenotype than evident from their independent effects^17,18^. Furthermore, prior studies to assess combinatorial effects have been inherently biased due to their need to minimize the search space by restricting the analysis to only a subset of genes chosen based on *a priori* knowledge^41–43^. Here, we provide a proof-of-concept analytical framework that remains agnostic to prior evidence and performs exhaustive searches to identify combinatorial effects among rare variants while retaining high granularity of data and interpretability of results.

We use our framework to identify gene pairs and triplets significantly associated with intellectual disability and show that several constituent genes are associated with nervous system processes. These mutated gene combinations are more likely to be inherited maternally or occur *de novo*, are depleted in unaffected siblings from the same family, and are less likely to involve canonical autism or ID genes, suggesting that genes forming significant combinations are less deleterious on their own but manifest effects only when combined with other similar genes carrying rare mutations. While previous studies have linked aggregate rare variant burden towards intellectual disability^44,45^, our results fine map the association to specific combinations of constituent genes contributing to the burden. We propose a novel paradigm for dissecting the complexity of genetic disorders, where an affected individual carries multiple combinations of rare variants, and each combination contributes to either the same phenotype or distinct phenotypes at varying effect sizes (**Figure 5**). A limitation of our method is that it tends to be biased towards genes that are mutated frequently enough to be observed in a combination. This limitation can be addressed by fixing specific primary variants of interest irrespective of their frequency and screening for “second-hit” modifiers that significantly co-occur with the primary variant, such as the co-occurrence of *RBM8A* variants in distal 1q21.1 deletion carriers manifesting thrombocytopenia-absent-radius syndrome and *TBX6* variants in 16p11.2 deletion carriers with scoliosis^46,47^.

**Figure 5:**
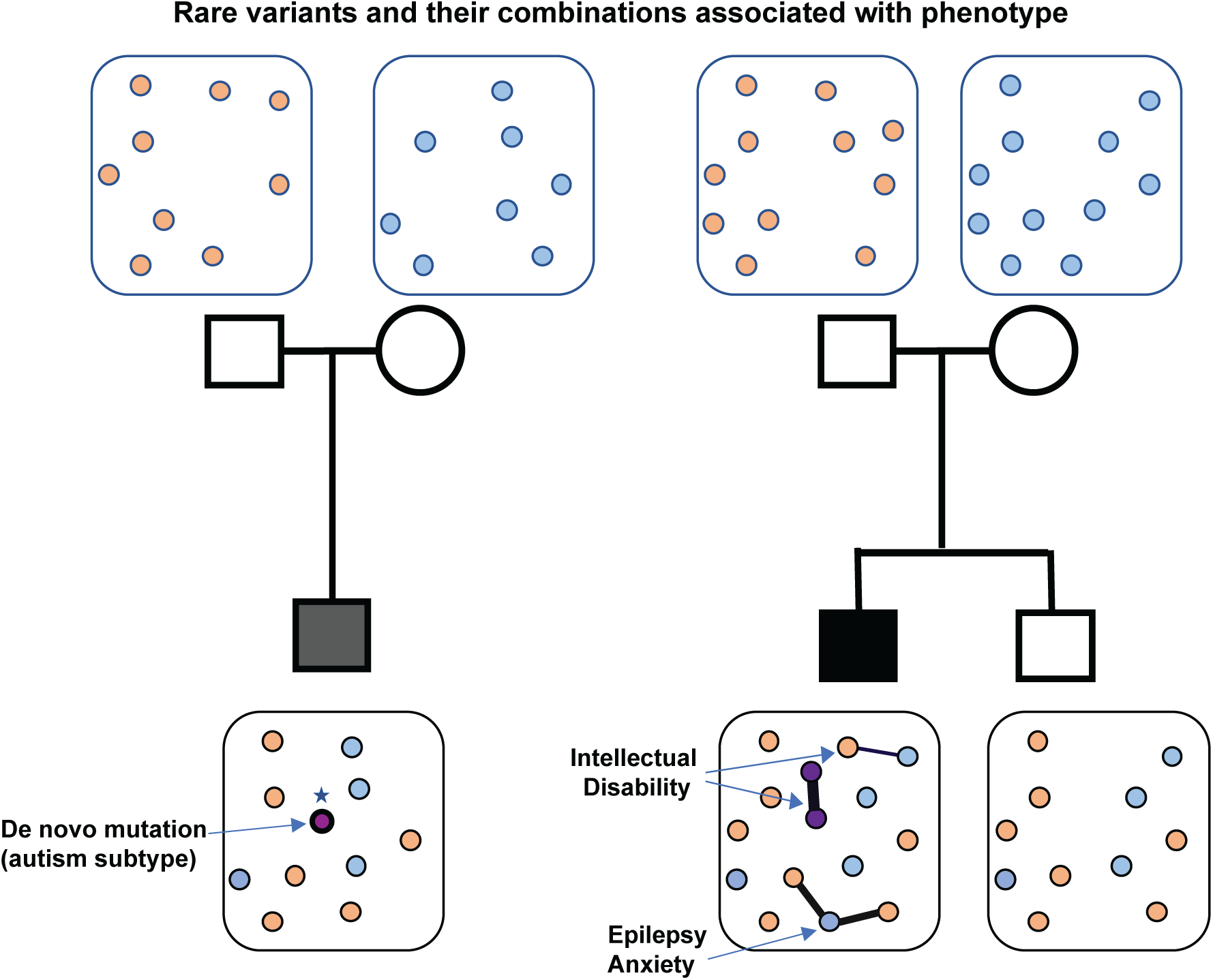
Rare variant models for complex disorders. The schematic shows two models for the genetic etiology of complex disorders. Circles represent rare variants present that are either *de novo* or inherited from a parent. On the left, individual high-effect *de novo* variants are strongly associated with a phenotype of interest. On the right, rare variants within an individual combine in multiple ways and contribute towards distinct phenotypes. The thickness of the connecting lines denotes effect sizes, and an affected individual can carry multiple oligogenic combinations of rare variants, each of which contributes to the same or distinct phenotypes. This extension of the oligogenic model enables further dissection of the genetic architecture of complex disorders.

Our method is fast and scalable, allows for fine-tuning combinatorial searches based on frequency, statistical power, and multiple testing criteria, and can be adapted to enable computational approximations to further improve run time and assess higher-order combinations beyond triplets. While larger sample sizes are generally required for detecting smaller frequency differences, we note that our framework achieves reliable statistical power even with modest sample sizes, implying that our framework could be applied to exome sequencing studies of other neurodevelopmental disorders that have not been explored for combinatorial effects. This approach can also be used to address a variety of research questions involving rare event combinations, including searching for protective effects of rare variants where simultaneous mutations are enriched in controls but not in cases, and finding combinations that exhibit specific enrichment or depletion patterns in more than two phenotypic groups. In summary, we provide a conceptual framework and the necessary tools to identify the oligogenic basis for complex disorders such as autism and intellectual disability, which hitherto was restricted to the analysis of canonical disorders such as Hirschsprung disease^48^ and Bardet-Biedl syndrome^12^.

## MATERIALS AND METHODS

We developed *RareComb* to address computational and statistical challenges associated with combinatorial analysis of rare variants. *RareComb* first uses the apriori algorithm to efficiently count the frequencies of co-occurring variant combinations. It then uses one-tailed binomial tests to compare the observed frequency of each variant combination to the expected frequency derived under the assumption of independence among the constituent variants within each combination (**Figure 1**). This method can be applied to identify variant combinations that are significantly enriched in cases but not in controls. In studies involving multiple comorbid phenotypes, this method can also be used to detect associations between specific combinations of variants and one or more (comorbid) phenotypes (see **Supplementary Note**). The general principles of our method, built using the basic axioms of probability theory, can be easily extended to a variety of problems involving rare higher-order combinations (**Supp. Figure 11**).

### Identifying frequencies of rare variant combinations

*RareComb* utilizes the apriori algorithm to efficiently calculate frequencies of variant combinations from sparse Boolean matrices (of 0s and 1s) (**Supp. Figure 12A**). The apriori algorithm has been successfully applied to analyze consumer behavior, where identifying products frequently purchased together could benefit a company^49,50^. While an algorithm that is used to derive insights from patterns within highly frequent events (i.e. frequent itemset mining) might not seem like a good fit to analyze rare variant combinations, its ability to perform disciplined search based on both built-in and user-specified constraints makes it an ideal counting tool. *For example*, the apriori algorithm avoids enumerating each of the 50 million pairs or 167 billion triplets from just 10,000 variants, and instead prunes the search-space based on user-defined criteria such as minimum frequency threshold and size of combinations (pairs, triplets, etc.) (**Supp. Figure 12B**). *RareComb* applies an additional constraint to the algorithm to limit its search to co-occurring events, which further reduces the search space (see **Supplementary Note)**. *For example*, when considering variants A and B, only the frequency of the presence of both variants (A=1 & B=1) is counted, and not absence of either or both variants (A=1 & B=0; A=0 & B=1; or A=0 & B=0).

### Statistical Inference

*RareComb* utilizes the p-values of one-tailed binomial tests to establish the magnitude of enrichment for each rare variant combination (**Figure 1**). For each combination, *RareComb* formulates null and alternate hypotheses for the binomial test by considering the event of observing all constituent variants together within a group of individuals as success and all other possibilities as failure in a binomial trial:

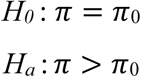

where,

*π* = Probability of *observing* all constituent rare variants of a combination together within a cohort, i.e., P(A=1 & B=1)

*π*_0_ = *Expected* probability derived from the frequency of individual variants of a combination, under the assumption of independence, i.e., P(A=1) * P(B=1).

*RareComb* then compares the null binomial distribution derived using the sample size of the group (n) and the expected probability (*π*_0_) (i.e., X ∼ Binom(n, p = *π*_0_)) with the observed probability (*π*), and calculates the probability of observing rare variants occurring together at least as frequently as they were observed within the cohort (i.e. p-value).

In case-control analyses, this method is applied independently to each group, and the p-values between them are compared. The combinations exhibiting enrichment in both cases and controls, likely due to proximity of variants in linkage disequilibrium, are eliminated, following which the p-values in cases are adjusted for multiple-testing to identify statistically significant combinations that exhibit enrichment in cases but not in controls. Finally, the effect sizes are calculated using Cohen’s d and the statistical power is measured using 2-sample 2-proportion tests, as additional metrics to prioritize the final set of significant rare variant combinations. In genotype-comorbid phenotype association analyses, the method is applied just once to the entire cohort, with multiple-testing adjusted p-values serving as a sufficient metric to identify high quality associations between genotypes and two or more co-occurring phenotypes.

### Statistical power and computational performance of the method

We measured the relationship between sample size and statistical power for both binomial and 2-sample 2-proportion tests used in the framework. It took 1,356 samples for the binomial test to achieve a statistical power of 80% to establish statistical enrichment between expected and observed co-occurrence frequencies of 0.1% and 0.5% (**Supp. Figure 13**). This number increased to 6,469 when the test needed to be more sensitive to compare frequencies of 0.3% and 0.5%. Similarly, it took 7,840 samples for the 2-sample 2-proportion test to achieve 80% power to establish statistical difference between co-occurrence frequencies of 2% and 0.5% observed in two groups (**Supp. Figure 14**). The sample size requirement increased to 14,633 to differentiate frequencies of 1.5% and 0.5% at 80% statistical power. These results align with the known relationship between sample size and statistical power, and indicate that our method can be reliably applied to analyze reasonably modest-size cohorts.

We also measured the run times for the case-control analysis to identify significant pairs and triplets of mutated genes using simulated data of three discrete sizes of samples (5,000, 10,000, and 50,000 individuals) and genes (5,000, 10,000, and 15,000 genes). The apriori algorithm was run on single-core CPUs with 256 GB memory and was constrained to analyze combinations observed in at least 0.15% of the samples. Given the memory-intensive nature of the apriori algorithm implemented in the ‘arules’ package, 256 GB was chosen to maintain uniformity^51^. However, smaller input files could be processed successfully using much less memory. As expected, the runtimes were proportional to the size of the combination (pairs versus triplets) and the number of input variables (**Supp. Figure 15**). While the increase in run time with the increase in sample size is apparent for pairs, lower runtimes observed with running 50,000 samples compared to 5,000 samples for triplets can be attributed to stochasticity of the input data. Overall, the analysis of gene pairs took between one minute and 12 minutes while triplets took between two minutes and 150 minutes. Since several factors influence the runtime of the method, a trial-and-error approach to determine an optimal minimum frequency threshold for co-occurring events can help identify relevant combinations without resulting in insufficient memory due to combinatorial explosion.

### Samples

We used whole exome sequencing data from 6,189 affected males from the Simons Foundation Powering Autism Research (SPARK)^21^ and 1,878 affected males from 2,247 simplex families from the Simons Simplex Collection (SSC)^52^ cohort from the Simons Foundation Autism Research Initiative (SFARI)^53^. We selected only male probands for our analysis to avoid any confounding effect due to gender or ascertainment bias^54,55^. While diagnosis information for intellectual disability (ID), anxiety, attention deficit hyperactivity disorders (ADHD), schizophrenia, language and sleep disorders were encoded as binary variables for the SPARK samples, full-scale intelligence quotient (IQ) scores were available for the SSC cohort.

### Data preparation and quality control

Variant Call Format (VCF) files obtained from exome sequencing data were annotated using ANNOVAR^56^ for rsID information and variant frequency using ExAC^57^ and gnomAD^58^. To overcome the limitations of using a single method to predict pathogenicity, the effects of non-synonymous mutations were annotated using 11 prediction methods: SIFT^59^, Polyphen2^60^ (HDIV), Polyphen2 (HVAR), LRT^61^, MutationTaster^62^, MutationAssessor^63^, FATHMM^64^, MetaSVM^65^, PROVEAN^66^, REVEL^67^, and CADD^68^. Briefly, all missense, stop-loss/gain, and start-loss/gain variants within exonic, 3’, and 5’ UTR regions with minor allele frequencies ≤1% identified based on both ExAC and gnomAD databases were selected. Then, variants with allele depth of ≥15 and allele balance between 25% and 75% for heterozygous variants and > 90% for homozygous variants were selected as high-quality variants. Deleteriousness of the variants were measured and reported differently by each prediction method. REVEL provided a score between 0 and 1, with higher scores indicating higher level of deleteriousness, while Polyphen2 and MutationAssessor classified variants into one of three categories. *For example*, Polyphen2 classified variants as ‘Deleterious’, ‘Possibly damaging’, or ‘Tolerated’, while MutationAssessor classified variants as ‘High’, ‘Medium’, or ‘Low’. The other nine methods classified variants as either ‘Deleterious’ or ‘Tolerated’. Pathogenicity reported by each tool was encoded as a binary variable, with the categories ‘Possibly damaging’ and ‘Medium’ encoded as 0.5. Thus, the composite pathogenicity score derived from the 10 tools could range between 0 and 10. Missense variants with a cumulative score of ≥4 and stop-loss/gain predicted as ‘deleterious’ either based on CADD score (CADD phred >30) or MutationTaster were considered deleterious for all analyses. Indels and other smaller structural variants were not considered, as their functional impact could not be easily assessed.

### Gene Ontology (GO) and Human Phenotype Ontology (HPO) enrichment analyses

Gene Ontology term enrichment analyses were performed using the ‘Gene Ontology API’ accessed using the ‘post’ command of the python package ‘requests’ (python version 3.7)^31^. All analyses were performed using parameters for *homo sapiens* (organism = ‘9606’) to identify biological processes enrichment (annotDataSet = ‘GO:0008150’) using binomial tests. HPO enrichment analyses were performed using data from the ‘genes_to_phenotype’ file obtained from the HPO website^32^. Since enrichment of phenotypes is not automatically evaluated by HPO, we used customized R scripts to derive baseline expectations that could be compared against the actual observations to determine significance using the p-values from binomial tests.

### Statistical analysis

All statistical analyses were performed using R v3.6.1 (R Foundation for Statistical Computing, Vienna, Austria)^69^ and Python (v3.7)^70^. All data-related plots were generated using the R package ggplot2^71^.

### Software Availability

*RareComb* is available as an open-source (https://github.com/girirajanlab/RareComb) R package that can be downloaded from the Comprehensive R Archive Network (CRAN) repository^72^. It can also be installed into development environments via interfaces such as Rstudio^73^ using the command *install*.*packages(‘RareComb’)*. The tool provides several functionalities that allow users to run the types of analyses described in this manuscript. The functionalities are as follows: (1) Identify rare event combinations statistically enriched within a single group; (2) Identify rare event combinations statistically enriched in cases but not in controls; (3) Identify rare event combinations enriched in cases but depleted in controls; (4) Identify statistically enriched rare event combinations that include at least one element from an user-supplied list; and (5) Identify genotypes statistically enriched within individuals manifesting two or more comorbid phenotypes. Each functionality takes a Boolean matrix as input and provides a set of user-adjustable parameters to customize the analysis, and delivers the results in a tabular format as csv files. Detailed instructions on the available functionalities and parameters built into *RareComb* and their usage can be found on the GitHub page or CRAN website. A shiny app illustrating the ideas behind *RareComb* is available online at https://girirajanlab.shinyapps.io/RareComb/ ^74^.

## DECLARATIONS

### Ethics approval and consent to participate

As these data were de-identified, all our samples were exempt from IRB review and conformed to the Helsinki Declaration. No other approvals were needed for the study.

### Consent for publication

All authors agree and consent for publication of the manuscript.

### Authors’ contributions

VK and SG conceived the project. VK performed the analyses, generated the plots/images, and wrote and revised the manuscript; SG supervised the research and wrote and revised the manuscript. All authors read and approved the final draft of the manuscript.

## Acknowledgements

We thank Naomi Altman, Yifei Huang, Dajiang Liu, Matthew Jensen, and Corrine Smolen for constructive comments on the manuscript. This work was supported by R01-MH107431, R01-GM121907, Seed Grants program from the Institute of Computational and Data Sciences at Penn State, and resources from the Huck Institutes of the Life Sciences (to SG). The funding bodies had no role in data collection, analysis, and interpretation. The authors are grateful to all the families who participated in the SSC and SPARK consortia, as well as the principal investigators, clinical sites, and staff for the consortia. The authors appreciate obtaining access to genetic and phenotypic data for SPARK and SSC through the Simons Foundation Autism Research Initiative (SFARI) Base. Approved researchers can obtain the SSC and SPARK population datasets described in this study by applying at https://base.sfari.org.

## References

1. Coventry, A. et al. Deep resequencing reveals excess rare recent variants consistent with explosive population growth. Nat. Commun. 1, 131 (2010).

2. Keinan, A. & Clark, A. G. Recent explosive human population growth has resulted in an excess of rare genetic variants. Science. 336, 740–743 (2012).

3. Tennessen, J. A. et al. Evolution and functional impact of rare coding variation from deep sequencing of human exomes. Science. 337, 64–69 (2012).

4. McClellan, J. & King, M. C. Genetic heterogeneity in human disease. Cell 141, 210–217 (2010).

5. Wilfert, A. B. et al. Recent ultra-rare inherited variants implicate new autism candidate risk genes. Nat. Genet. 53, 1125–1134 (2021).

6. Iossifov, I. et al. The contribution of de novo coding mutations to autism spectrum disorder. Nature 515, 216–221 (2014).

7. Sebat, J. et al. Strong Association of De Novo Copy Number Mutations with Autism. Science. 316, 445–449 (2007).

8. Badano, J. L. & Katsanis, N. Beyond mendel: An evolving view of human genetic disease transmission. Nat. Rev. Genet. 3, 779–789 (2002).

9. Gifford, C. et al. Oligogenic inheritance of a human heart disease involving a genetic modifier. Science. 364, 865–870 (2019).

10. Pizzo, L. et al. Rare variants in the genetic background modulate cognitive and developmental phenotypes in individuals carrying disease-associated variants. Genet. Med. 21, 816–825 (2019).

11. Girirajan, S. et al. A recurrent 16p12.1 microdeletion supports a two-hit model for severe developmental delay. Nat. Genet. 42, 203–209 (2010).

12. Badano, J. L. et al. Dissection of epistasis in oligogenic Bardet-Biedl syndrome. Nature 439, 326–330 (2006).

13. Leblond, C. S. et al. Genetic and functional analyses of SHANK2 mutations suggest a multiple hit model of autism spectrum disorders. PLoS Genet. 8, e1002521 (2012).

14. Uricchio, L. H., Zaitlen, N. A., Ye, C. J., Witte, J. S. & Hernandez, R. D. Selection and explosive growth alter genetic architecture and hamper the detection of causal rare variants. Genome Res. 26, 863–873 (2016).

15. Halvorsen, M. et al. Increased burden of ultra-rare structural variants localizing to boundaries of topologically associated domains in schizophrenia. Nat. Commun. 11, 1842 (2020).

16. Krumm, N. et al. Excess of rare, inherited truncating mutations in autism. Nat. Genet. 47, 582–588 (2015).

17. Murdoch, W. J., Singh, C., Kumbier, K., Abbasi-Asl, R. & Yu, B. Definitions, methods, and applications in interpretable machine learning. Proc. Natl. Acad. Sci. U. S. A. 116, 22071–22080 (2019).

18. Molnar, C., Casalicchio, G. & Bischl, B. Interpretable Machine Learning – A Brief History, State-of-the-Art and Challenges. Commun. Comput. Inf. Sci. 1323, 417–431 (2020).

19. Agarwal, R. & Srikant, R. Fast algorithms for mining association rules. Proc. 20th VLDB Conf.

20. Agrawal, Rakesh; Ramakriahnan, S. Fast Algorithms for Mining Association Rules. Proc. 20th VLDB Conf. 487–499 (1994).

21. The SPARK Consortium. SPARK: A US Cohort of 50,000 Families to Accelerate Autism Research. Neuron 97, 488–493 (2018).

22. Fischbach, G. D. & Lord, C. The simons simplex collection: A resource for identification of autism genetic risk factors. Neuron 68, 192–195 (2010).

23. Girirajan, S. et al. Phenotypic Heterogeneity of Genomic Disorders and Rare Copy-Number Variants. N. Engl. J. Med. 367, 1321–1331 (2012).

24. Turner, T. N. et al. Genomic Patterns of De Novo Mutation in Simplex Autism. Cell 171, 710–722 (2017).

25. Dodé, C. et al. Loss-of-function mutations in FGFR1 cause autosomal dominant Kallmann syndrome. Nat. Genet. 33, 463–465 (2003).

26. Schell, U. et al. Mutations in FGFR1 and FGFR2 cause familial and sporadic pfeiffer syndrome. Hum. Mol. Genet. 4, 323–328 (1995).

27. Dazzo, E. et al. Heterozygous Reelin Mutations Cause Autosomal-Dominant Lateral Temporal Epilepsy. Am. J. Hum. Genet. 96, 992–1000 (2015).

28. Yoshinaga, T. et al. A novel frameshift mutation of SYNE1 in a Japanese family with autosomal recessive cerebellar ataxia type 8. Hum. Genome Var. 4, 17052 (2017).

29. Synofzik, M. et al. SYNE1 ataxia is a common recessive ataxia with major non-cerebellar features: A large multi-centre study. Brain 139, 1378–1393 (2016).

30. Prasad, A. et al. A Discovery resource of rare copy number variations in individuals with autism spectrum disorder. G3 Genes, Genomes, Genet. 2, 1665–1685 (2012).

31. Mi, H., Muruganujan, A., Ebert, D., Huang, X. & Thomas, D. PANTHER version 14 : more genomes, a new PANTHER GO-slim and improvements in enrichment analysis tools. Nucleic Acids Res. 47, 419–426 (2019).

32. Köhler, S. et al. The human phenotype ontology in 2021. Nucleic Acids Res. 49, D1207–D1217 (2021).

33. Gilger, J. W., Pennington, B. F. & DeFries, J. C. A Twin Study of the Etiology of Comorbidity: Attention-deficit Hyperactivity Disorder and Dyslexia. J. Am. Acad. Child Adolesc. Psychiatry 31, 343–348 (1992).

34. Krumm, N. et al. Transmission disequilibrium of small CNVs in simplex autism. Am. J. Hum. Genet. 93, 595–606 (2013).

35. Guo, H. et al. Genome-wide copy number variation analysis in a Chinese autism spectrum disorder cohort. Sci. Rep. 7, 44155 (2017).

36. Guemez-Gamboa, A. et al. Inactivating mutations in MFSD2A, required for omega-3 fatty acid transport in brain, cause a lethal microcephaly syndrome. Nat. Genet. 47, 809–813 (2015).

37. Sanders, A. R. et al. Haplotypic association spanning the 22q11.21 genes COMT and ARVCF with schizophrenia. Mol. Psychiatry 10, 353–365 (2005).

38. Kenny, E. M. et al. Excess of rare novel loss-of-function variants in synaptic genes in schizophrenia and autism spectrum disorders. Mol. Psychiatry 19, 872–879 (2014).

39. Zheng, G. X. Y. et al. Haplotyping germline and cancer genomes with high-throughput linked-read sequencing. Nat. Biotechnol. 34, 303–311 (2016).

40. Girirajan, S. et al. Relative burden of large CNVs on a range of neurodevelopmental phenotypes. PLoS Genet. 7, e1002334 (2011).

41. Papadimitriou, S. et al. Predicting disease-causing variant combinations. Proc. Natl. Acad. Sci. U. S. A. 116, 11878–11887 (2019).

42. Kerner, G. et al. A genome-wide case-only test for the detection of digenic inheritance in human exomes. Proc. Natl. Acad. Sci. U. S. A. 117, 19367–19375 (2020).

43. Schaaf, C. P. et al. Oligogenic heterozygosity in individuals with high-functioning autism spectrum disorders. Hum. Mol. Genet. 20, 3366–3375 (2011).

44. Singh, T. et al. The contribution of rare variants to risk of schizophrenia in individuals with and without intellectual disability. Nat. Genet. 49, 1167–1173 (2017).

45. Fitzgerald, T. W. et al. Large-scale discovery of novel genetic causes of developmental disorders. Nature 519, 223–228 (2015).

46. Albers, C. A. et al. Compound inheritance of a low-frequency regulatory SNP and a rare null mutation in exon-junction complex subunit RBM8A causes TAR syndrome. Nat. Genet. 44, 435–439 (2012).

47. Yang, N. et al. TBX6 compound inheritance leads to congenital vertebral malformations in humans and mice. Hum. Mol. Genet. 28, 539–547 (2019).

48. Gabriel, S. B. et al. Segregation at three loci explains familial and population risk in Hirschsprung disease. Nat. Genet. 31, 89–93 (2002).

49. Brijs, T., Swinnen, G., Vanhoof, K. & Wets, G. Using association rules for product assortment decisions. ACM (1999). doi:10.1145/312129.312241

50. Glance, N. et al. Deriving marketing intelligence from online discussion. Proc. ACM SIGKDD Int. Conf. Knowl. Discov. Data Min. 419–428 (2005). doi:10.1145/1081870.1081919

51. Hahsler, M., Grun, B. & Hornik, K. arules – A Computational Environment for Mining Association Rules and Frequent Item Sets. J. Stat. Softw. 14, 1–6 (2005).

52. Sanders, S. J. et al. Insights into Autism Spectrum Disorder Genomic Architecture and Biology from 71 Risk Loci. Neuron 87, 1215–1233 (2015).

53. SFARI Base. https://www.sfari.org/resource/sfari-base/

54. Jacquemont, S. et al. A higher mutational burden in females supports a ‘female protective model’ in neurodevelopmental disorders. Am. J. Hum. Genet. 94, 415–425 (2014).

55. Polyak, A., Rosenfeld, J. A. & Girirajan, S. An assessment of sex bias in neurodevelopmental disorders. Genome Med. 7, 94 (2015).

56. Wang, K., Li, M. & Hakonarson, H. ANNOVAR: Functional annotation of genetic variants from high-throughput sequencing data. Nucleic Acids Res. 38, e164 (2010).

57. Exome aggregation Consortium. Analysis of protein-coding genetic variation in 60,706 humans. Nature 536, 285–291 (2016).

58. Genome Aggregation Database Consortium. The mutational constraint spectrum quantified from variation in 141,456 humans. Nature 581, 434–443 (2020).

59. Ng, P. C. & Henikoff, S. SIFT: Predicting amino acid changes that affect protein function. Nucleic Acids Res. 31, 3812–4 (2003).

60. Adzhubei, I. A. et al. A method and server for predicting damaging missense mutations. Nat. Methods 7, 248–249 (2010).

61. Chun, S. & Fay, J. C. Identification of deleterious mutations within three human genomes. Genome Res. 19, 1553–1561 (2009).

62. Schwarz, J. M., Rödelsperger, C., Schuelke, M. & Seelow, D. MutationTaster evaluates disease-causing potential of sequence alterations. Nat. Methods 7, 575–576 (2010).

63. Reva, B., Antipin, Y. & Sander, C. Predicting the functional impact of protein mutations: Application to cancer genomics. Nucleic Acids Res. 39, 37–43 (2011).

64. Shihab, H. A. et al. Predicting the Functional, Molecular, and Phenotypic Consequences of Amino Acid Substitutions using Hidden Markov Models. Hum. Mutat. 34, 57–65 (2013).

65. Kim, S., Jhong, J. H., Lee, J. & Koo, J. Y. Meta-analytic support vector machine for integrating multiple omics data. BioData Min. 10, 2 (2017).

66. Choi, Y. & Chan, A. P. PROVEAN web server: A tool to predict the functional effect of amino acid substitutions and indels. Bioinformatics 31, 2745–2747 (2015).

67. Ioannidis, N. M. et al. REVEL: An Ensemble Method for Predicting the Pathogenicity of Rare Missense Variants. Am. J. Hum. Genet. 99, 877–885 (2016).

68. Rentzsch, P., Witten, D., Cooper, G. M., Shendure, J. & Kircher, M. CADD: Predicting the deleteriousness of variants throughout the human genome. Nucleic Acids Res. 47, D886–D894 (2019).

69. R Core Team. R: A language and environment for statistical computing. (2019).

70. Van Rossum, Guido; Drake, F. L. Python 3 Reference Manual. (2009).

71. Wickham, H. ggplot2: Elegant Graphics for Data Analysis. (Springer-Verlag New York, 2016).

72. CRAN Repository. Available at: https://cran.r-project.org/web/packages/available_packages_by_name.html.

73. RStudio Team. RStudio: Integrated Development for R. (2020).

74. Chang, W., Cheng, J., Allaire, J., Xie, Y. & McPherson, J. shiny: Web Application Framework for R. (2020).

