## Supplementary information for "Oligogenic combinations of rare variants influence specific phenotypes in complex disorders"

### A general framework for identifying rare variant combinations in complex disorders

### SUPPLEMENTAL TEXT

#### Definitions

- **Combination:** Multiple genomic entities considered together. Combinations with two entities constitute a pair, three entities constitute a triplet, and so on. *For example*, genes A, B and C can form four combinations: three pairs AB, AC and BC, and one triplet ABC.
- **Size/length of a combination:** Number of genomic entities under consideration. Pairs are of length 2 and triplets are of length 3.
- **Events:** Genomic events such as a structural variant or loss-of-function (LoF) or missense mutation observed within a single genomic unit such as a gene. Occurrence of an event is denoted as {Gene A = 1}.
- **Non-events:** Absence of genomic events within a given genomic unit, denoted as {Gene B = 0}.
- **Simultaneous events:** When events are observed in all constituent entities of a combination. Simultaneous occurrence of mutations in gene A and gene B are denoted as {Gene A=1 & Gene B=1}.
- **Simultaneous non-events:** When no events are observed in all constituent entities of a combination, denoted as {Gene A=0 & Gene B=0}.
- **Non-simultaneous events:** Events occur in at least one but not all constituent entities of a combination, denoted as {Gene A=1 & Gene B=0}, {Gene A=0 & Gene B=1 & Gene C=1}, etc.

#### A primer to the apriori algorithm

##### Combinatorial complexity

Current approaches for analysis of rare variants in complex disease deal with data sparsity by comparing aggregate enrichment of a specific variant (such as a CNV) or collective burden of variants between cases and controls. Analysis of combinations of rare events is challenging because substantially larger sample sizes are required to observe such events. *For example*, two independent rare variants 'rv1' and 'rv2' with a minor allele frequency of 5% (1 in 20 individuals) can be expected to be observed together only in 1/400 individuals. In fact, for every '*n*' samples required to observe a rare variant at a given allele frequency in a cohort, it takes at least $n^2$ samples to observe two rare variants of similar allele frequencies together. Both $n$ and $n^2$ increase exponentially with decreases in allele frequency thresholds. Even when large cohorts are available, an efficient algorithm is required to overcome the combinatorial explosion and efficiently calculate the frequency of simultaneously occurring rare events from sparse datasets.

##### List of parameters to constrain search and prune search space

The apriori algorithm is a breadth-first search algorithm that has become synonymous with two data mining techniques, '*association rule learning*' and '*frequent itemset mining*'. It is used to either automatically identify interesting associations between two or more variables called '*rules'*, of the format '{Gene A=1, Gene B=1} => {Phenotype=Severe}', or simply list frequently occurring set of items called 'itemsets', of the format {Gene A=1, Gene B=1, Gene C=1}, that meet a minimum frequency threshold (support) supplied to constrain the algorithm. Rules have two sides, the '*antecedent'* on the left and the '*consequent'* on the right, connected by the directionality of their association. The number of items in a rule is its *length*. The algorithm can report one or more items in the antecedent, but its consequent can only be a single item. Frequent itemsets are simply list of items without any relationship or directionality among them. The algorithm reports the itemsets along with their absolute frequencies. *For example*, the rule {Gene A=1, Gene B=1} => {Phenotype=Severe} has a length of 3, where {Gene A=1, Gene B=1} is the antecedent and {Phenotype=Severe} is the consequent. Similarly, {Gene A=1, Gene B=1, Gene C=1, Gene D=1} is an itemset of length 4.

Searching for patterns involving more than two items is computationally challenging due to the resulting combinatorial explosion. *For example*, to identify rules of length 4 using just 100 items, as many as 4 million possibilities ($100C_4$) must be considered, which sharply increases to 75 million if the length is increased to 5 ($100C_5$). The apriori algorithm addresses this challenge by constraining the search space using three important parameters that control the length, support/frequency, and confidence of the final set of rules in the output (**Supp. Figure 12**). *Confidence* is only applicable to rules, whereas the other two metrics are applicable to both rules and frequent itemsets. The algorithm systematically prunes the search space by using a subset of rules/itemsets of smaller lengths that meet the user-provided criteria for the three parameters to expand the search to rules/itemsets of larger lengths. This approach allows it to perform computationally efficient searches.

The three parameters used to constrain the algorithm are as follows:

1) *Length of the rule/itemset*: The length of the rule/itemset is the primary determinant of the search space due to its combinatorial relationship with the number of input items. While an upper limit for length is often supplied to the algorithm, a lower limit can also be specified if necessary.
2) *Support for the rule/itemset*: Support indicates the frequency in which the constituent items of an itemset or a rule appear together within a set of observations. If 20 out of 100 individuals carry the variants A and B together, then the support for the itemset {A=1, B=1} is 0.20. If all 20 individuals are associated with a 'severe' phenotype, then the support for the rule {A=1, B=1}=>{severe} is also 0.2. Each observation containing these three items serves as additional evidence supporting the existence of an association between the antecedent and the consequent. Providing a threshold for 'support' (lower limit) limits the search to only those itemsets and rules in which the constituent items appear together at least as frequently as the threshold.
3) *Confidence of the rule*: Confidence indicates the probability of observing the consequent when the antecedent is observed. For example, let's assume the antecedent {A=1, B=1} is observed in 25 out of 100 individuals (support for the antecedent = 0.25), and the antecedent {A=1, B=1} is observed together with the consequent {severe phenotype} in 20 of those instances (i.e., support for the rule {A=1, B=1}=>{severe phenotype} is 0.20). Here, 80% of all individuals carrying variants A and B together have a severe phenotype, meaning that confidence = [support for the rule/support for the antecedent] = 0.2/0.25 = 80%. Since rules with high confidence could be predictive of the outcome, a lower limit for this parameter is often provided to the algorithm.

It should be noted that the confidence metric reported for a rule does not take the frequency of the *consequent* within the cohort into account. *For example*, if there are three times as many cases as controls in a cohort (for a binary outcome), a genotype combination that occurs three times as frequently in cases than controls would have a confidence of 75%. i.e., $3/4^{th}$ of all co-occurring genotypic events is associated with one of the two possible outcomes. While 75% confidence might suggest that a genotype combination is predictive of the outcome, it was achieved simply due to the relatively higher frequency of one of the two binary outcomes in the cohort. A fourth useful parameter named '*Lift'* takes this limitation into consideration and adjusts the confidence by controlling for the frequency of the consequent, where Lift = [Confidence/Frequency of the consequent]. So, the lift for an antecedent 'A' and the consequent 'C' is [P(A∩C)/P(A)]/P(C) or simply P(A∩C)/P(A)* P(C). Basic axioms of probability dictate that if A and C are independent, P(A∩C) would be the same as P(A)*P(C), making *lift* a measure of the extent of dependence between the antecedent and the consequent. Conversely, if the *lift* score is 1, the antecedent and consequent are independent of each other. Therefore, the *lift* score is directly proportional to the dependence between the *antecedent* and the *consequent*. After incorporating the frequency of the consequent in the prior example, the *lift* score becomes 1. The *lift* score can hence be used to identify truly dependent events among all high confidence rules. Unlike length, support, and confidence, thresholds for *lift* cannot be supplied to constrain the apriori algorithm prior to invoking it, but can instead be used *post-hoc* to identify high quality rules.

#### **Using the apriori algorithm**

The ability of the apriori algorithm to generate results in two formats, '*frequent itemsets*' and '*rules*', provides flexibility to leverage it in multiple ways. These two formats allow genotypes to be either analyzed independently or together with the phenotypes. '*Frequent itemsets'* is the best fit for counting the frequency of simultaneous events involving only the genotypes. For analyses involving both genotypes and phenotypes, where any meaningful combination must involve at least a single phenotype, 'rules' are the best fit since the algorithm can be constrained to include only the phenotypes as the *consequent* item in the rule. We use both formats for counting the frequency of simultaneous events along with the general principles of statistical inference in two specific ways: one involving just the genotypes for case/control comparisons, and the other involving both genotypes and phenotypes for assessing comorbidities.

##### **Case/control enrichment analysis**

*RareComb* can be used to identify genotype combinations that exhibit differential enrichment of simultaneous events between two groups. As genotypes are analyzed exclusively in this approach, 'frequent itemsets' are generated using the apriori algorithm. The minimum frequency of simultaneous events (support threshold) in cases is provided as the initial constraint to the algorithm, while being agnostic to the absolute frequency in controls. Once all genotype combinations in which the frequency of simultaneous events is at least as high as the support threshold in cases are identified, the p-values from the binomial test quantifying the magnitude of enrichment relative to the expectation under the assumption of independence are calculated. For the subset of combinations with higher-than-expected frequency of simultaneous events in cases, the corresponding p-values in the control group are calculated. The combinations in which simultaneous events occur more frequently than expected in both cases and controls are discarded, since they signify potentially dependent genomic events (due to factors such as linkage disequilibrium). Finally, combinations that are statistically significant in cases after adjusting the p-values for multiple testing while remaining non-significant in controls are considered to impact disease/phenotype.

##### ***Comorbidity analysis***

Understanding the genetic basis of comorbid phenotypes associated with complex diseases is challenging due to two main reasons. *First*, when multiple phenotypes are considered, even in a large cohort, the number of individuals with a specific combination of phenotypes is very small. For every '*n*' binary phenotype, $2^n$ configurations of comorbidities are possible, and it is challenging to find adequate samples representing each configuration in complex disease cohorts. *Second*, even if the sample size is large enough to have adequate individuals for each configuration, current methods are not able to effectively explain or predict associations with combinations of phenotypes. In many analyses, either a new outcome variable is created based on the composite configuration of the phenotypes, or one is derived based on how the comorbid phenotypes cluster among themselves within the cohort. We extend our method to overcome existing limitations to provide explanations for multiple phenotypes considered together as individual units. While retaining granular phenotypes reduces the size of samples available within each configuration, the rarity of such phenotypic configurations can be turned into an advantage by screening for genotypes that are observed together more frequently than expected within such rare phenotype configurations. Our framework can be used to measure the likelihood of observing a set of phenotypes and genotypes together as frequently as they are observed within the cohort using the frequencies of individual items. *For example*, let's assume that 7 individuals in a cohort of size 2,000 are diagnosed with three phenotypes 'p1', 'p2' and 'p3' simultaneously, and 5 of them carry deleterious mutations simultaneously in genes 'g1' and 'g2'. Let's also assume that phenotypes p1, p2 and p3 and genotypes 'g1' and 'g2' are each observed independently in exactly 20 individuals within the cohort (1% of the cohort). One of the axioms of probability dictates that if these five events are independent, the probability of observing them all together in an individual is $(0.01)^5$, making the odds of observing the combination 'p1', 'p2', 'p3', 'g1', 'g2' in 5 individuals by chance alone extremely unlikely. The method applies this reasoning to identify combinations of phenotypes that occur together with combinations of genotypes more frequently than expected by chance alone. Such a method can be challenging due to the exorbitant number of combinations to be evaluated ($100C_5$ = ~75 million; $100C_5$ = ~255 billion), but we address this challenge using the apriori algorithm to search for combinations that occur at least as frequently as the support threshold provided to constrain the algorithm.

The method analyzes the entire cohort and generates combinations that meet the input criterion for '*support'* provided to the apriori algorithm. For this approach, the apriori algorithm is made to generate 'rules' with two specific constraints. *First*, only phenotypes are eligible to be the *consequent* item. *Second*, both genotypes and phenotypes are eligible to appear in the list of items in the *antecedent* portion of the rules. These constraints both limit the search space and ensure that any combination reported by the algorithm includes at least a single phenotype. Once the combinations that meet all input criteria for the apriori algorithm are obtained, the p-values from the binomial tests are calculated by comparing the expected frequency with the observed frequency of these qualifying combinations. The combinations that remain significant after multiple testing correction are identified as genotypes that contribute towards the comorbid phenotypes.

### SUPPLEMENTAL FIGURES

#### Technical workflow of *RareComb* to analyze for pairs of genes

| | G1 | G2 | G3 |
|---|---|---|---|
| Case 1 | 1 | 0 | 0 |
| Case 2 | 0 | 0 | 0 |
| Case 3 | 0 | 1 | 1 |
| Case 4 | 0 | 0 | 0 |
| Case 5 | 1 | 1 | 1 |
| Control 1 | 0 | 1 | 0 |
| Control 2 | 0 | 0 | 1 |
| Control 3 | 1 | 0 | 0 |
| Control 4 | 0 | 0 | 0 |
| Control 5 | 1 | 1 | 0 |

Step 1 | Split cases from controls

Step 2 | Calculate the frequency of individual events in each group

Step 3 | Run the apriori algorithm independently on each group to generate frequent itemsets for simultaneous events.

Item Size = 1

| Events | Support |
|---|---|
| {G1 = 1} | 2/5 |
| {G2 = 1} | 2/5 |
| {G3 = 1} | 2/5 |

Item Size = 2

| Event Pairs {X,Y} | Support |
|---|---|
| {G1 = 1, G2 = 1} | 1/5 |
| {G1 = 1, G3 = 1} | 1/5 |
| {G2 = 1, G3 = 1} | 2/5 |

Item Size = 1

| Events | Support |
|---|---|
| {G1 = 1} | 2/5 |
| {G2 = 1} | 2/5 |
| {G3 = 1} | 1/5 |

Item Size = 2

| Event Pairs {X,Y} | Support |
|---|---|
| {G1 = 1, G2 = 1} | 1/5 |
| {G1 = 1, G3 = 1} | 0 |
| {G2 = 1, G3 = 1} | 0 |

Step 4 | Obtain P(X ∩ Y)

| X | Y | X ∩ Y | Observed P(X ∩ Y) | Expected P(X ∩ Y) |
|---|---|---|---|---|
| G1 = 1 | G2 = 1 | {G1 = 1, G2 = 1} | 1/5 = 0.2 | 2/5 * 2/5 = 0.16 |
| G1 = 1 | G3 = 1 | {G1 = 1, G3 = 1} | 1/5 = 0.2 | 2/5 * 2/5 = 0.16 |
| G2 = 1 | G3 = 1 | {G2 = 1, G3 = 1} | 2/5 = 0.4 | 2/5 * 2/5 = 0.16 |

Step 4 | Obtain P(X ∩ Y)

| X | Y | X ∩ Y | Observed P(X ∩ Y) | Expected P(X ∩ Y) |
|---|---|---|---|---|
| G1 = 1 | G2 = 1 | {G1 = 1, G2 = 1} | 1/5 = 0.2 | 2/5 * 2/5 = 0.16 |
| G1 = 1 | G3 = 1 | {G1 = 1, G3 = 1} | 0 | 2/5 * 1/5 = 0.08 |
| G2 = 1 | G3 = 1 | {G2 = 1, G3 = 1} | 0 | 2/5 * 1/5 = 0.08 |

Step 5 | Run binomial tests

| X | Y | Observed | Expected | P(X> = x) \| p-value |
|---|---|---|---|---|
| G1 = 1 | G2 = 1 | 0.2 | 0.16 | 0.58 |
| G1 = 1 | G3 = 1 | 0.2 | 0.16 | 0.58 |
| G2 = 1 | G3 = 1 | 0.4 | 0.16 | 0.18 |

Step 5 | Run binomial tests

| X | Y | Observed | Expected | P(X> = x) \| p-value |
|---|---|---|---|---|
| G1 = 1 | G2 = 1 | 0.2 | 0.16 | 0.58 |
| G1 = 1 | G3 = 1 | 0 | 0.08 | 1.00 |
| G2 = 1 | G3 = 1 | 0 | 0.08 | 1.00 |

Step 6 | Adjust for multiple tests & compare p-values

| X | Y | Cases P(X> = x) \| p-value | Controls P(X> = x) \| p-value |
|---|---|---|---|
| G1 = 1 | G2 = 1 | 0.58 | 0.58 |
| G1 = 1 | G3 = 1 | 0.58 | 1.00 |
| G2 = 1 | G3 = 1 | 0.18 | 1.00 |

Step 7 | Identify combinations with complementary p-value trend

**Supp. Figure 1: Technical workflow of *RareComb*.** *RareComb* uses an input Boolean matrix consisting of variant information and binary phenotypic outcome for case-control analysis. It then applies the apriori algorithm independently to cases and controls to obtain the frequencies of simultaneously occurring events that meet the selection thresholds for length (pairs, triplets, etc.) and frequency. Binomial tests are applied to each eligible combination independently within cases and controls. Gene combinations are considered significant (after multiple-testing correction), when mutations are observed simultaneously in their constituent genes more frequently than expected under the assumption of independence among them, in cases but not in controls.

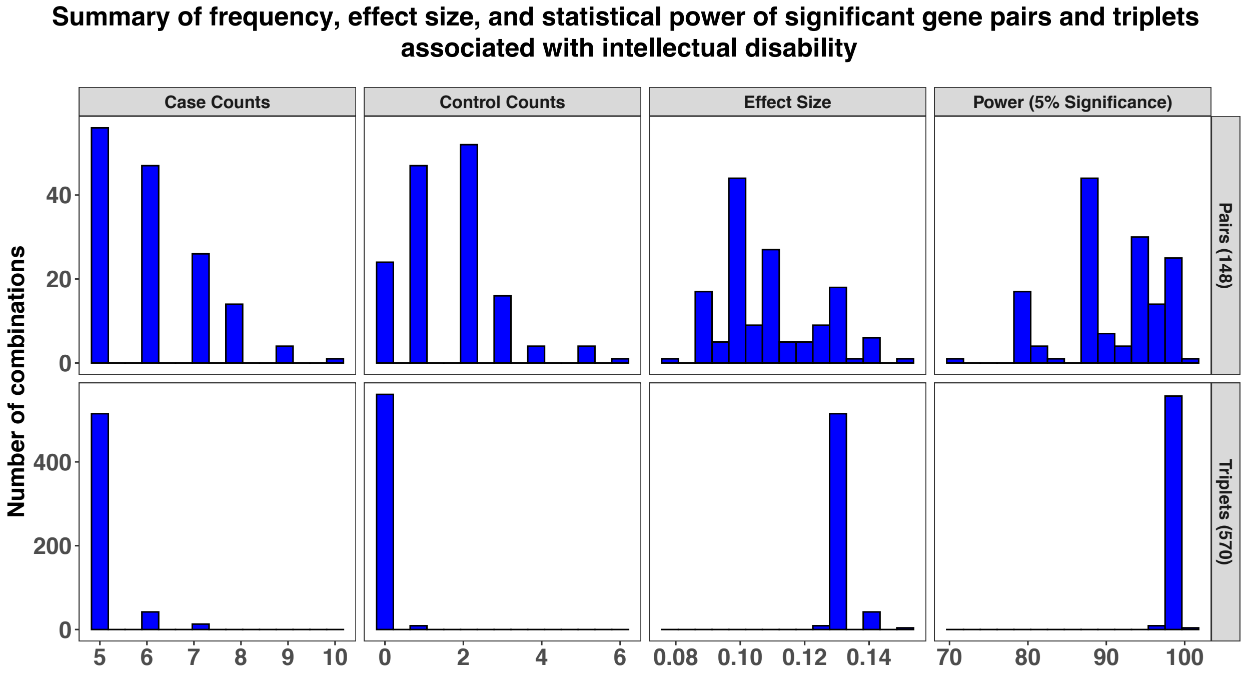

**Supp. Figure 2: Summary of significant gene pairs and triplets identified from the SPARK cohort.** Higher frequencies of simultaneous mutations are observed for pairs than for triplets (‘Case Counts’/’Control Counts’ panels along the X-axis), since simultaneous events tend to occur less frequently for combinations of larger sizes than smaller sizes. Notably, most significant triplets were observed in five cases, whereas significant pairs were observed in five or more number of cases. Effect sizes (Cohen’s d) quantify the differences in absolute frequency of combinations in cases versus controls. Since we used a statistical power cut-off >90% to identify 570 significant triplets, effect size and power are not comparable between pairs and triplets.

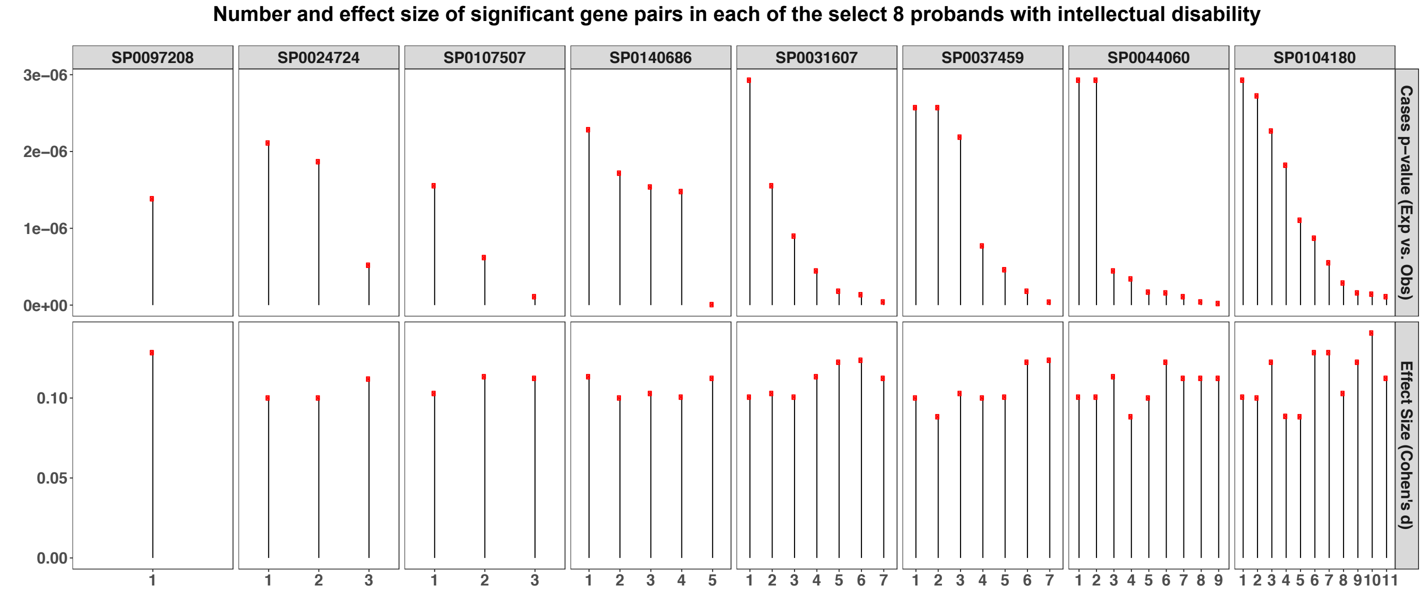

**Supp. Figure 3: The range of p-values and Cohen's d for mutated gene pairs in a representative set of probands from the SPARK cohort.** This figure illustrates that an individual can carry more than one combination of mutated genes significantly associated with the same phenotype, with each combination showing different enrichment (from binomial tests) and effect sizes (Cohen's d). Data from eight representative probands, each carrying multiple significant pairs of mutated genes, are shown here. The X-axis corresponds to probands, and the Y-axis shows p-values from binomial tests in cases and effect sizes measured using Cohen's d.

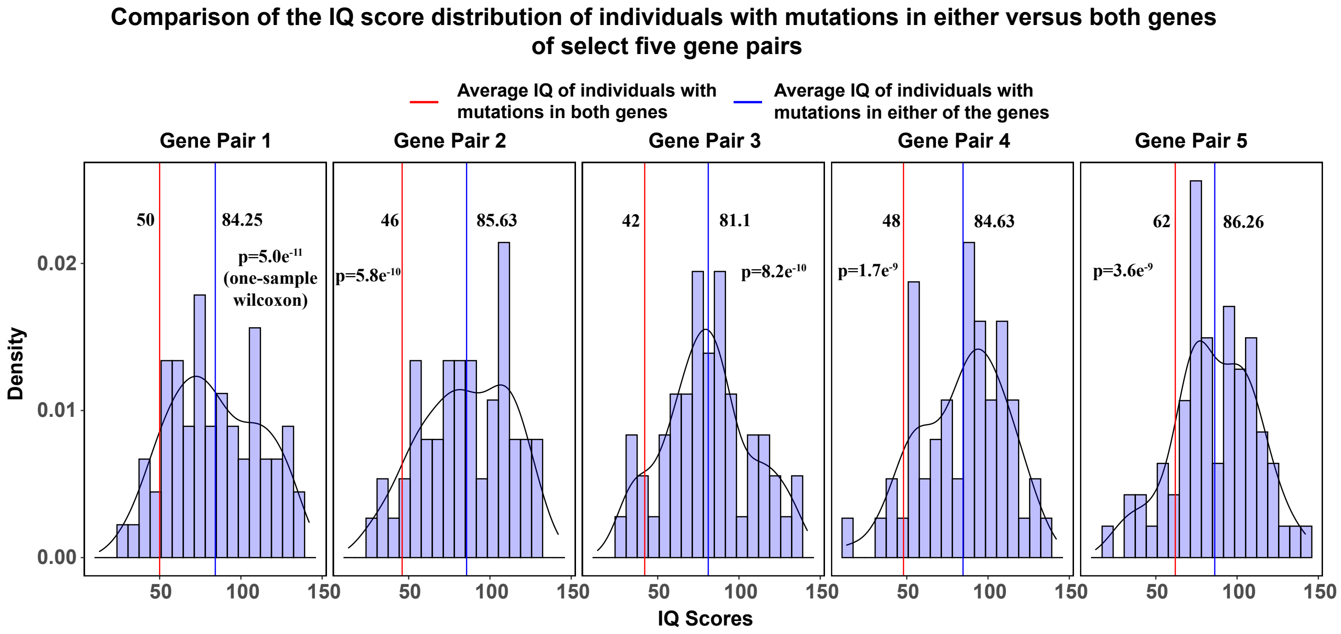

**Supp. Figure 4: Comparison of IQ scores of individuals carrying mutations in either of the constituent genes (SSC Cohort) with those carrying mutations in both genes of significant gene pairs.** Distributions of IQ scores from individuals carrying mutations in either of the two genes from select five mutated gene pairs are shown. The blue line indicates the mean of the distribution of IQ scores, and the red line indicates the mean IQ of individuals carrying mutations in both genes. We find that carriers of both mutations tend to have lower IQ scores on average compared to the average IQ of carriers of either of the two mutations.

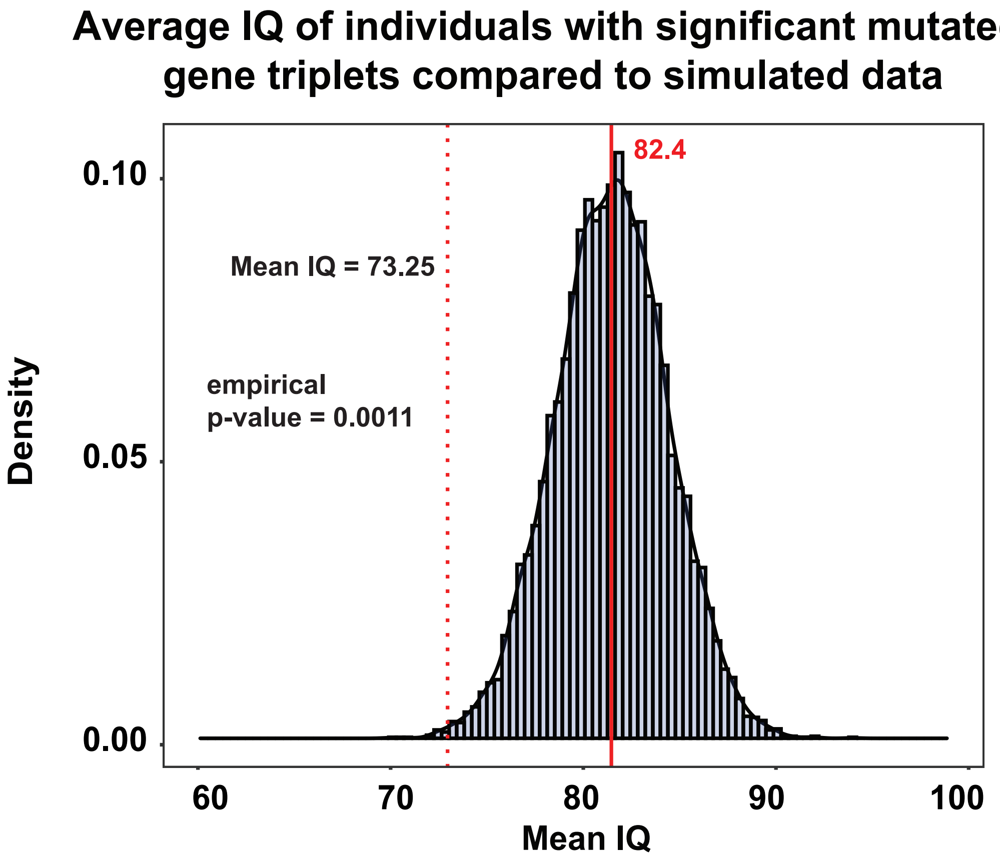

**Supp. Figure 5: Comparison of IQ scores of carriers of mutations in significant gene triplets compared with the simulated distribution**. The mean IQ score of individuals carrying significant gene triplets in the SSC cohort (73.25) is significantly lower (empirical p-value = 0.0013) when compared to the distribution of mean IQ scores (82.4) obtained from the simulation (see Figure 2A).

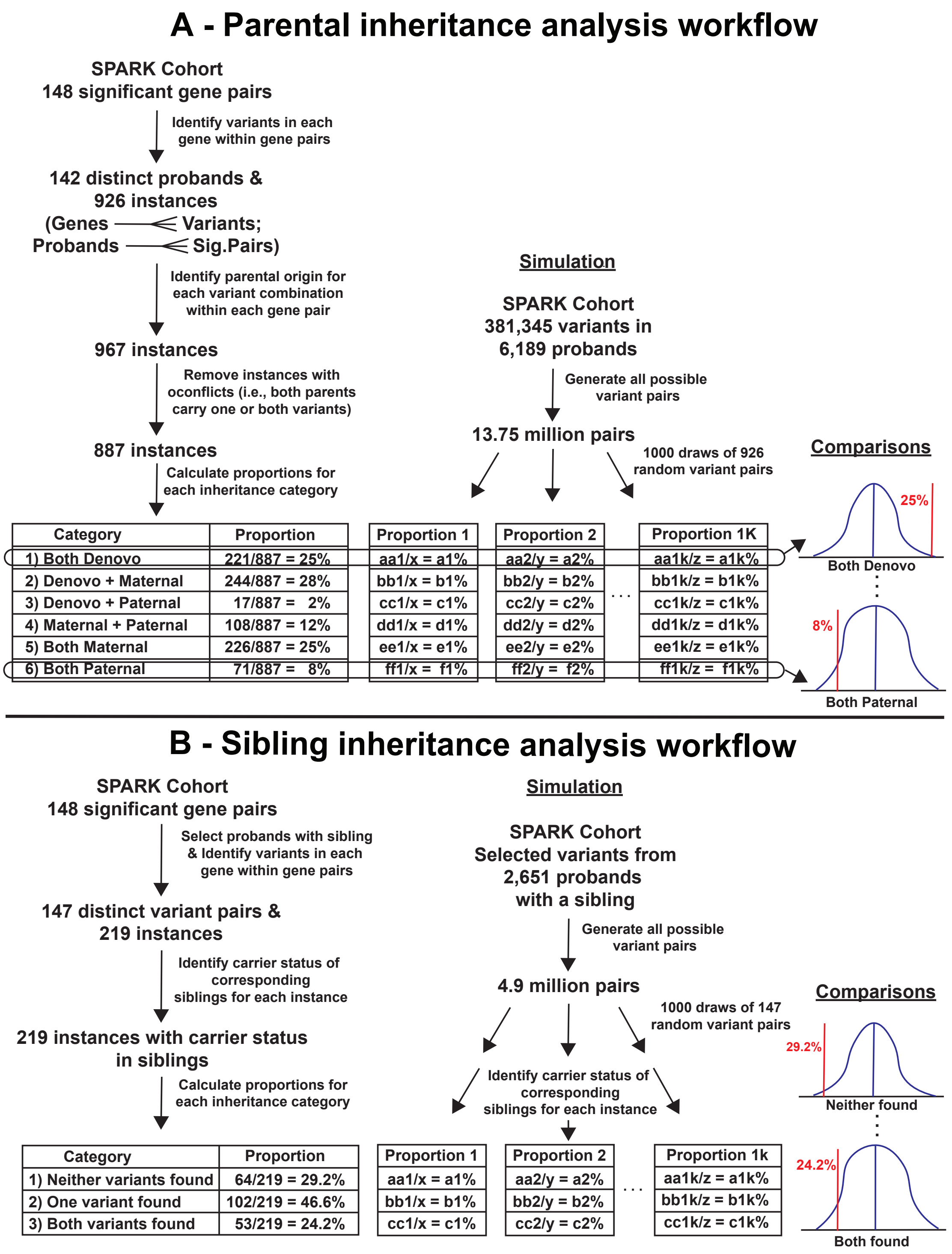

**Supp. Figure 6: Analysis of parental inheritance patterns.** (A) Outline of the steps involved in identifying the parental inheritance pattern of significant gene pairs and comparing them with distributions obtained from simulations. (B) Outline of the steps involved in identifying the carrier status of significant gene pairs in siblings and comparing them to distributions obtained from simulations.

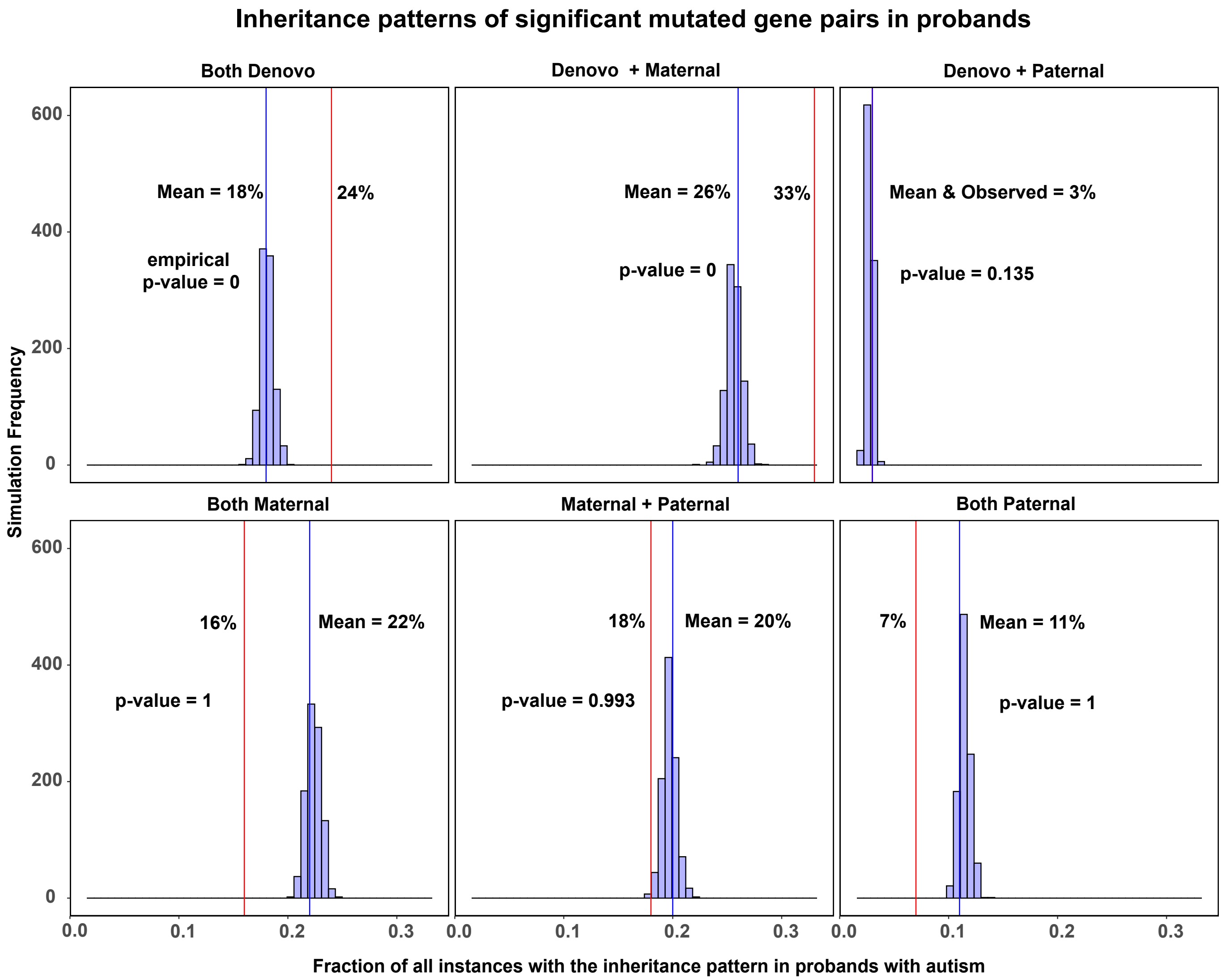

**Supp. Figure 7: Analysis of parental inheritance pattern of significant gene pairs associated with autism from the SPARK cohort.** Histograms show the fraction of all instances of mutated genes in a combination that belong to each of the six possible inheritance patterns compared to simulated distributions. Significant pairs were obtained by applying *RareComb* to SPARK data from probands as cases compared to parents as controls. For each simulation, the inheritance status of random pairs of mutated genes from the cohort were identified, and the fraction of those instances belonging to one of the six categories was calculated. Comparing the observed fractions with the mean of simulated fractions show statistically significant enrichment for instances when both variants are *de novo* or when one variant is *de novo* and the other transmitted from the mother.

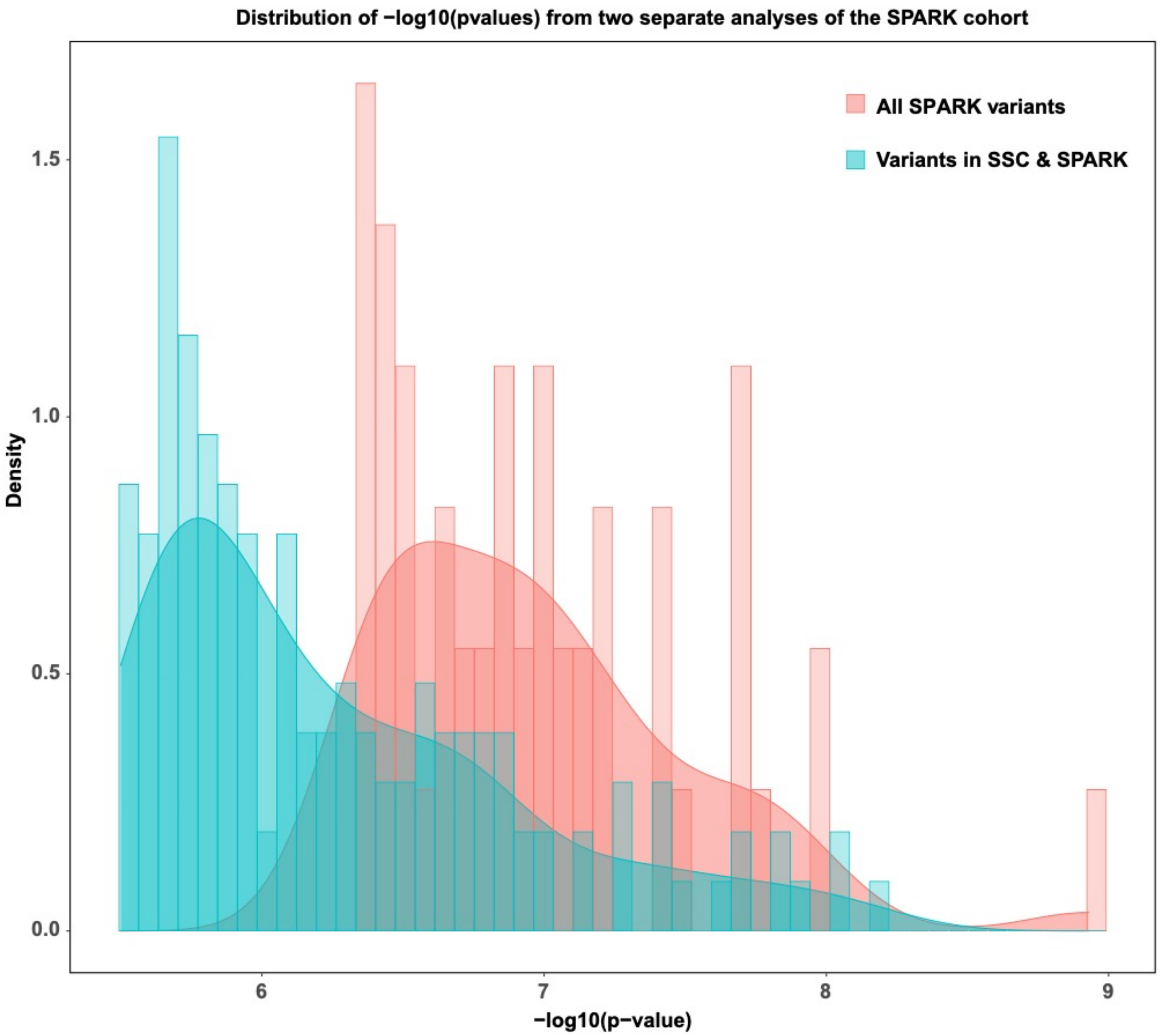

**Supp. Figure 8: Comparison of p-values between 52 (obtained using all SPARK variants) and 148 significant gene pairs (obtained using variants observed in both SPARK & SSC cohorts).** The shift in the distribution of p-values between the two analyses reflects the fact that combinations with *more* significant p-values could be observed when the method is applied to a larger set of genes compared to analysis using a smaller gene set. The larger the sample space of genes, the higher the likelihood of finding highly significant combinations.

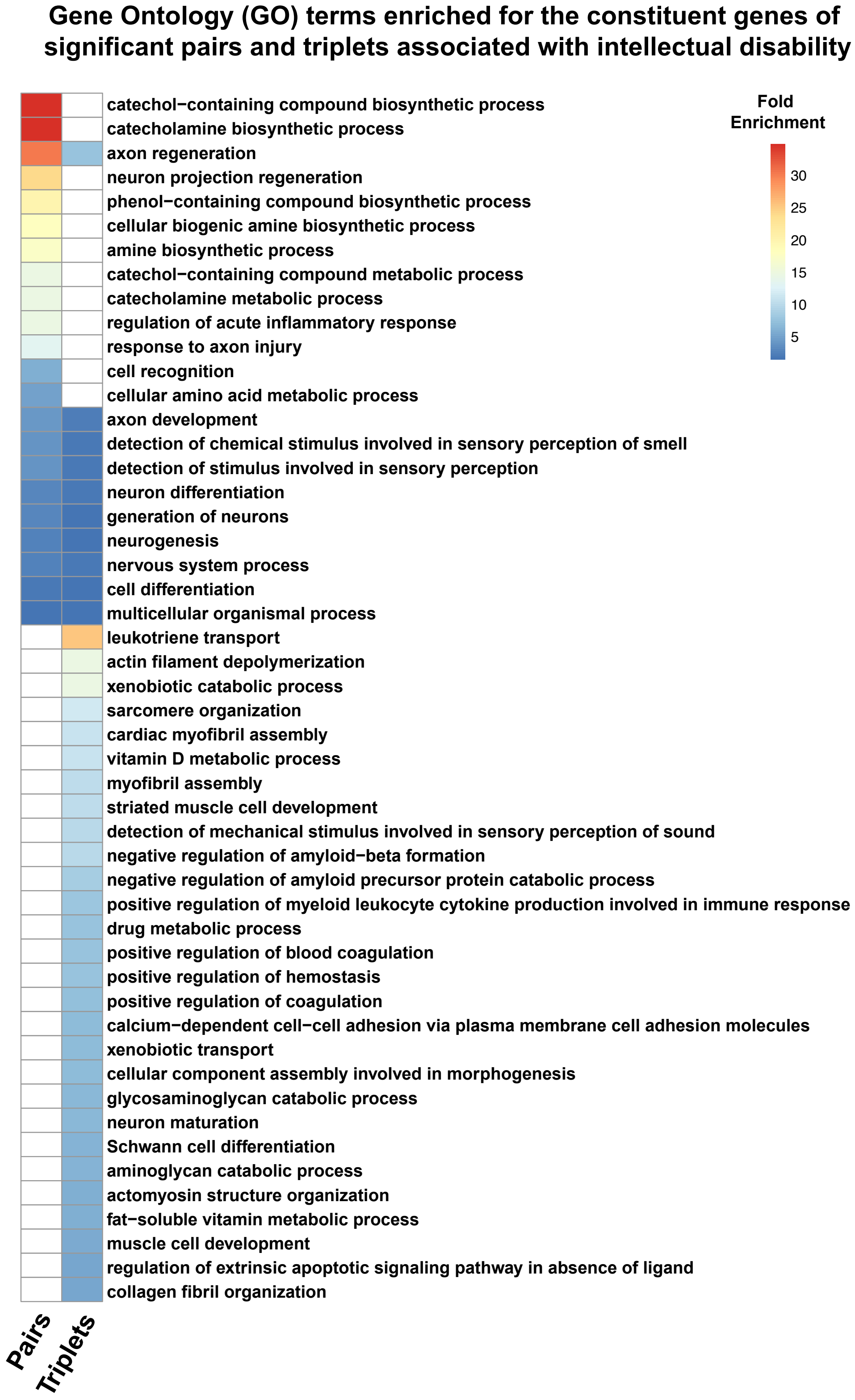

**Supp. Figure 9: GO term enrichment analysis for genes within significant pairs and triplets.** Fold enrichment of GO terms identified as statistically significant using the binomial test are listed. Seven of the nine enriched GO terms shared between the genes from significant pairs and triplets were associated with nervous system development and function. For example, several neurotransmitter-related terms showed as high as 40-fold enrichment for genes from the significant pairs.

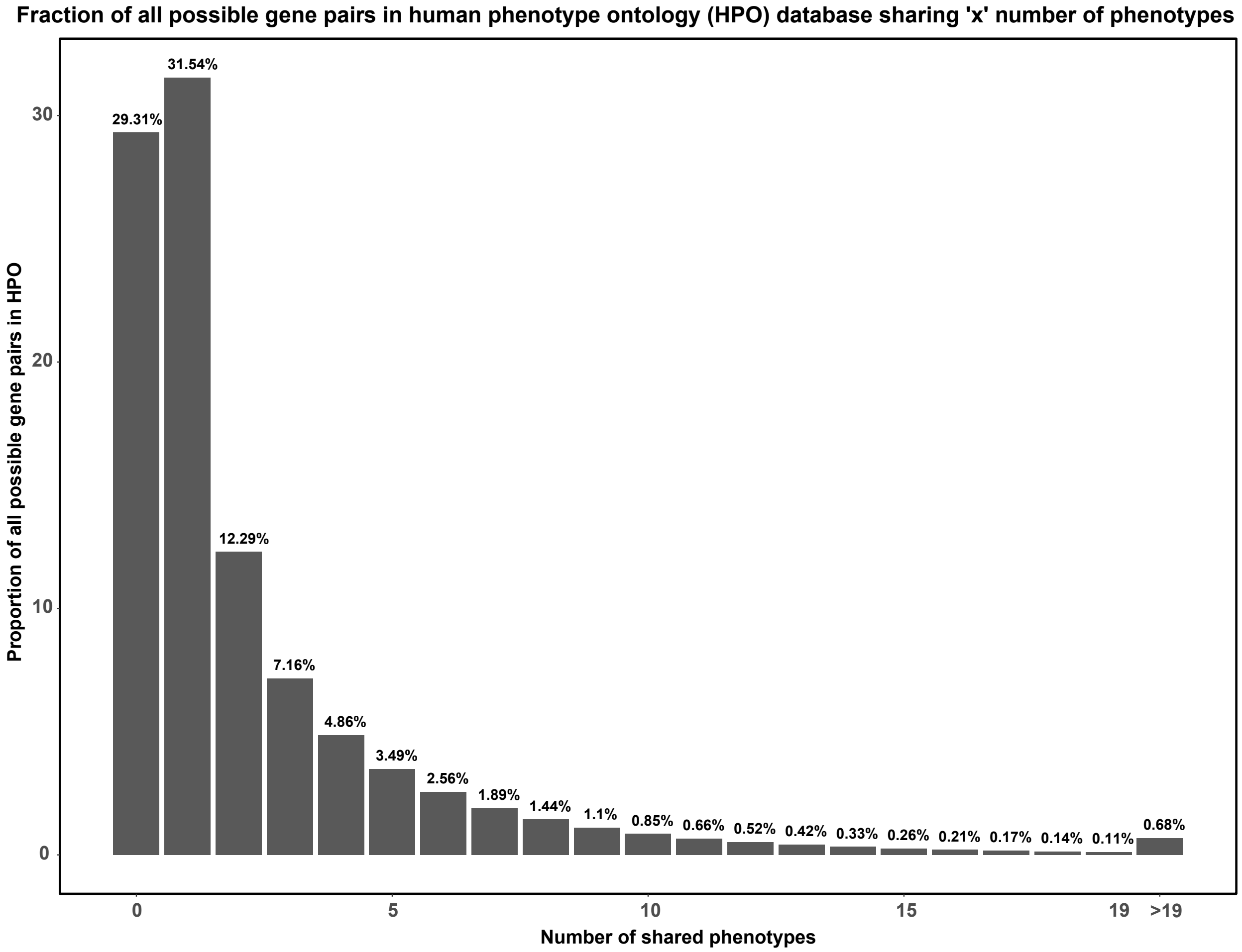

**Supp. Figure 10: Distribution of the expected number of phenotypes shared between two genes within HPO.** Barplot represents the number of phenotypes shared by each of the ~10 million gene pairs formed by 4,484 genes from HPO. We found that 60.9% of the pairs shared either no phenotype (29.31%) or a single phenotype (31.54%) with each other. These proportions serve as expected baselines for the binomial tests to compare and identify the significance of the number of phenotypes shared between the significant gene pairs identified by *RareComb*.

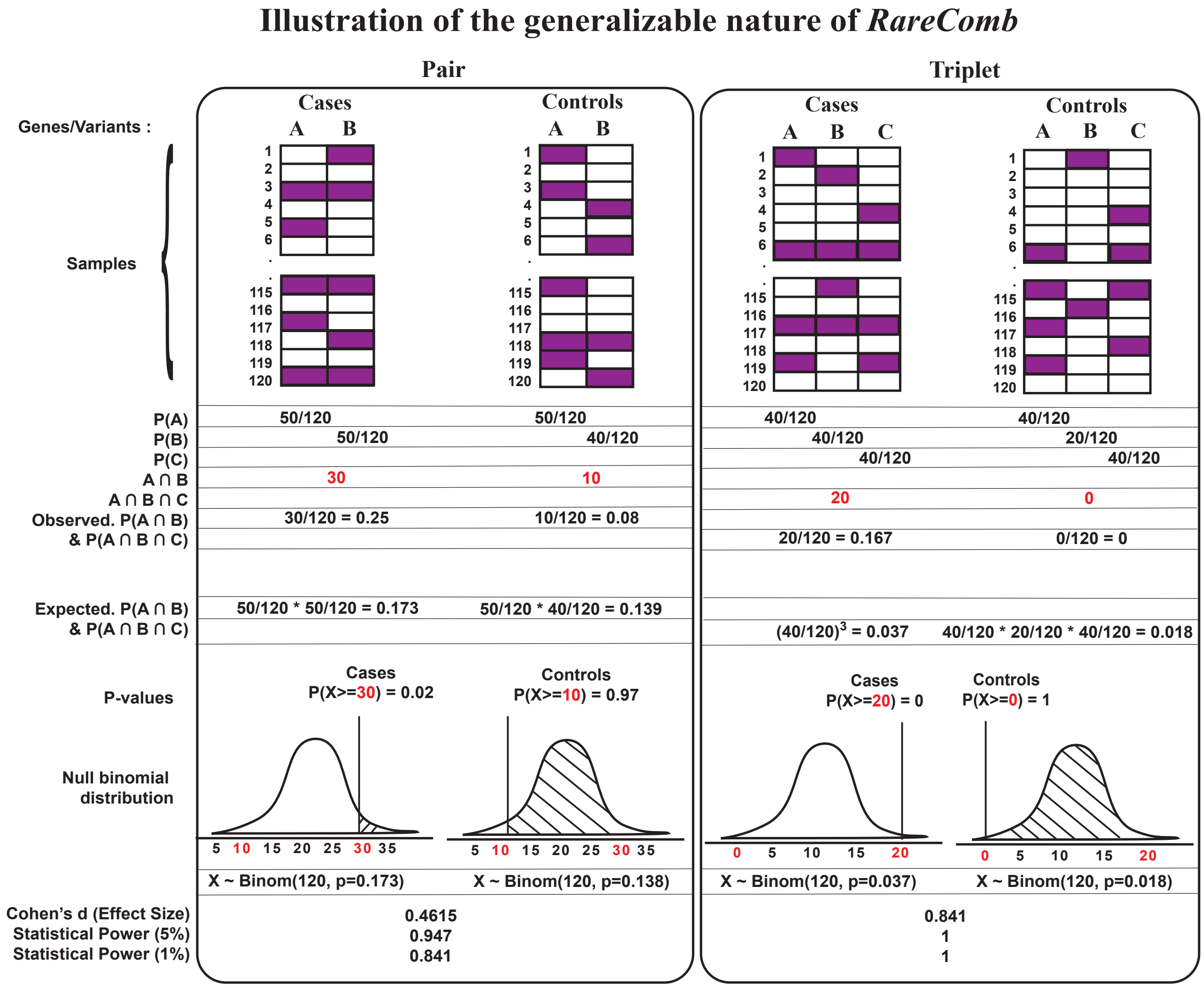

**Supp. Figure 11: Generalizable nature of *RareComb* illustrated using specific examples for pairs and triplets.** The principles of probability theory were used to derive the probability of co-occurring events expected under the assumption of independence for the constituent events. This principle was used to calculate significance of mutated gene pairs, and triplets, and can be extended to identify other higher-order combinations.

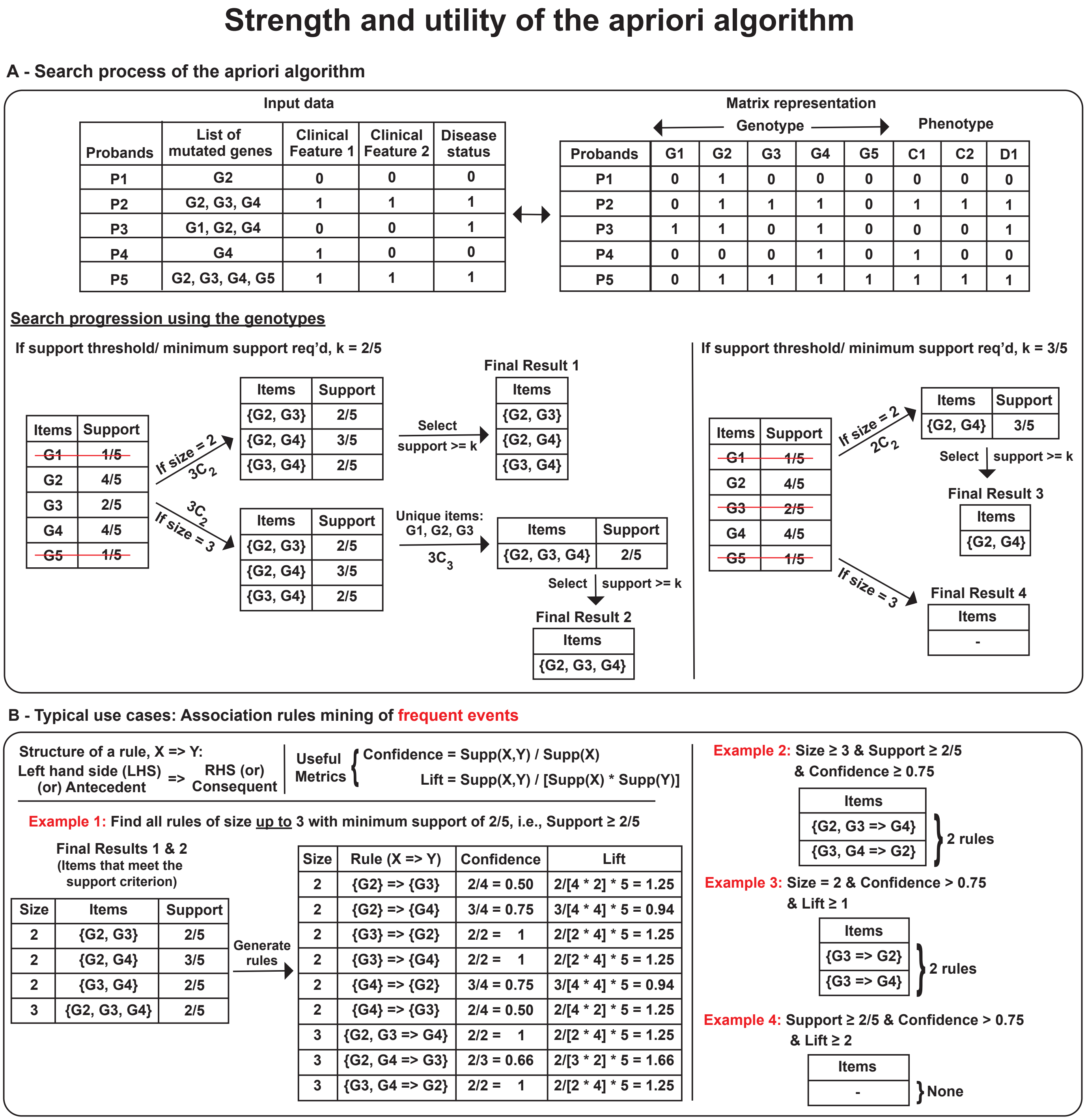

**Supp. Figure 12: A primer to the apriori algorithm and association rule mining. (A)** Diagram showing the search progression of the apriori algorithm. The apriori algorithm implemented in the R package 'arules' takes Boolean input data and searches for the frequency of simultaneous events efficiently by continually pruning the search space during each step of its progression, allowing it to enumerate the frequencies of combinations in a reasonable amount of time. **(B)** A typical application of the apriori algorithm is for association rule mining to identify interesting relationships among highly frequent events. Parameters such as length, support and confidence are used to both constrain the algorithm and to prioritize significant associations.

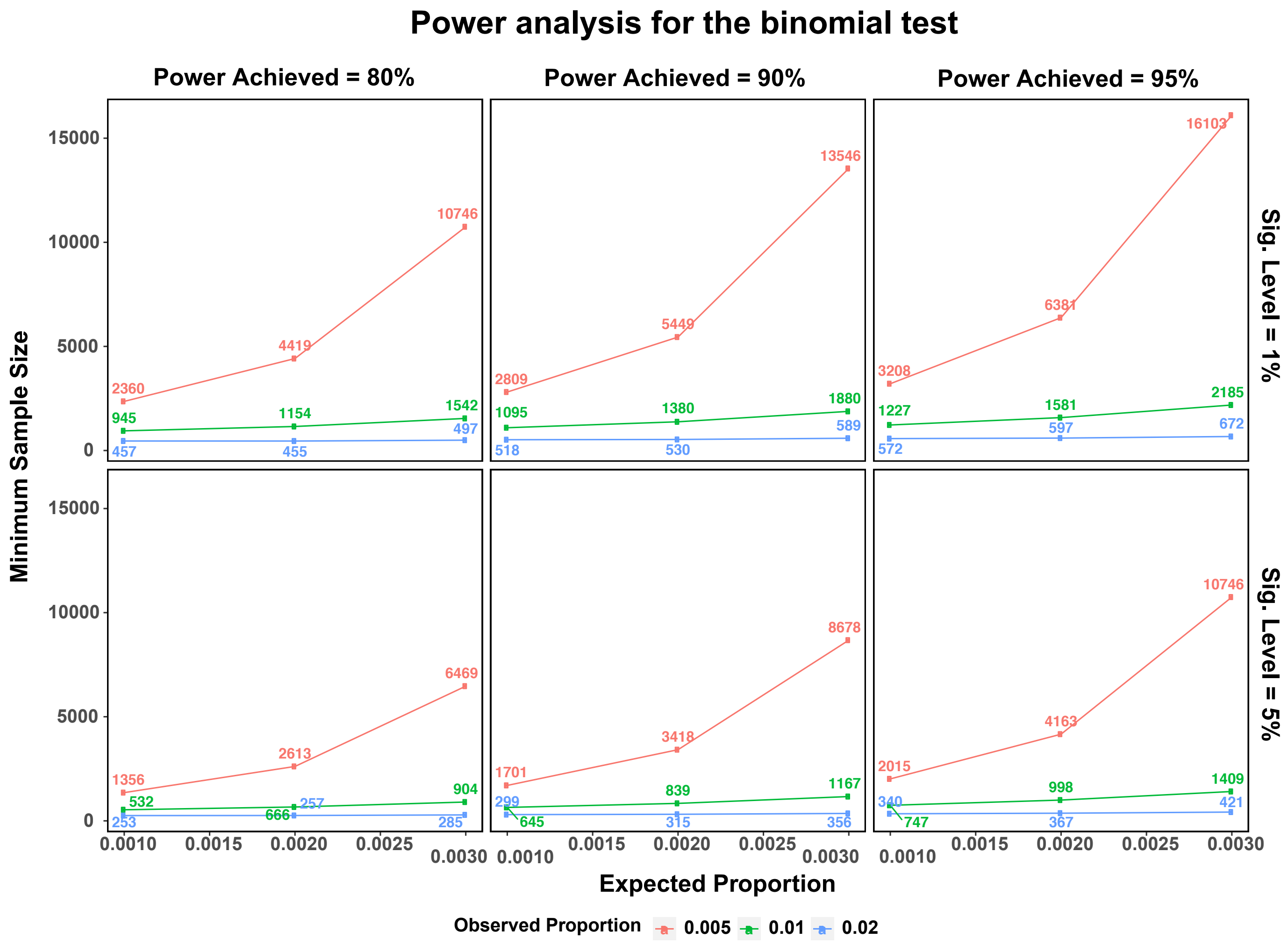

**Supp. Figure 13: Power analysis of binomial tests to compare expected versus observed frequencies of co-occurring events.** The panels along the X-axis show the minimum number of samples required for binomial tests to meet statistical powers of 80, 90 and 95% respectively, while the panels along the Y-axis show the sample size requirements at 1% and 5% statistical significance thresholds. Values along the X-axis represent the expected frequency of co-occurring events (0.1%, 0.2% and 0.3%) in cases, and line colors correspond to three specific frequencies (0.5%, 1% and 2%) in which co-occurring events are observed in cases. The results demonstrate that higher sample sizes are needed when comparisons must be sensitive enough to detect minor differences between expected and observed frequencies of co-occurring events, whereas relatively smaller sample sizes may be sufficient to achieve higher statistical power when such (i.e., exp. vs obs.) frequency differences are larger. Similarly, as expected, larger sample sizes are warranted for binomial tests to achieve high statistical power and to meet more stringent statistical significance thresholds.

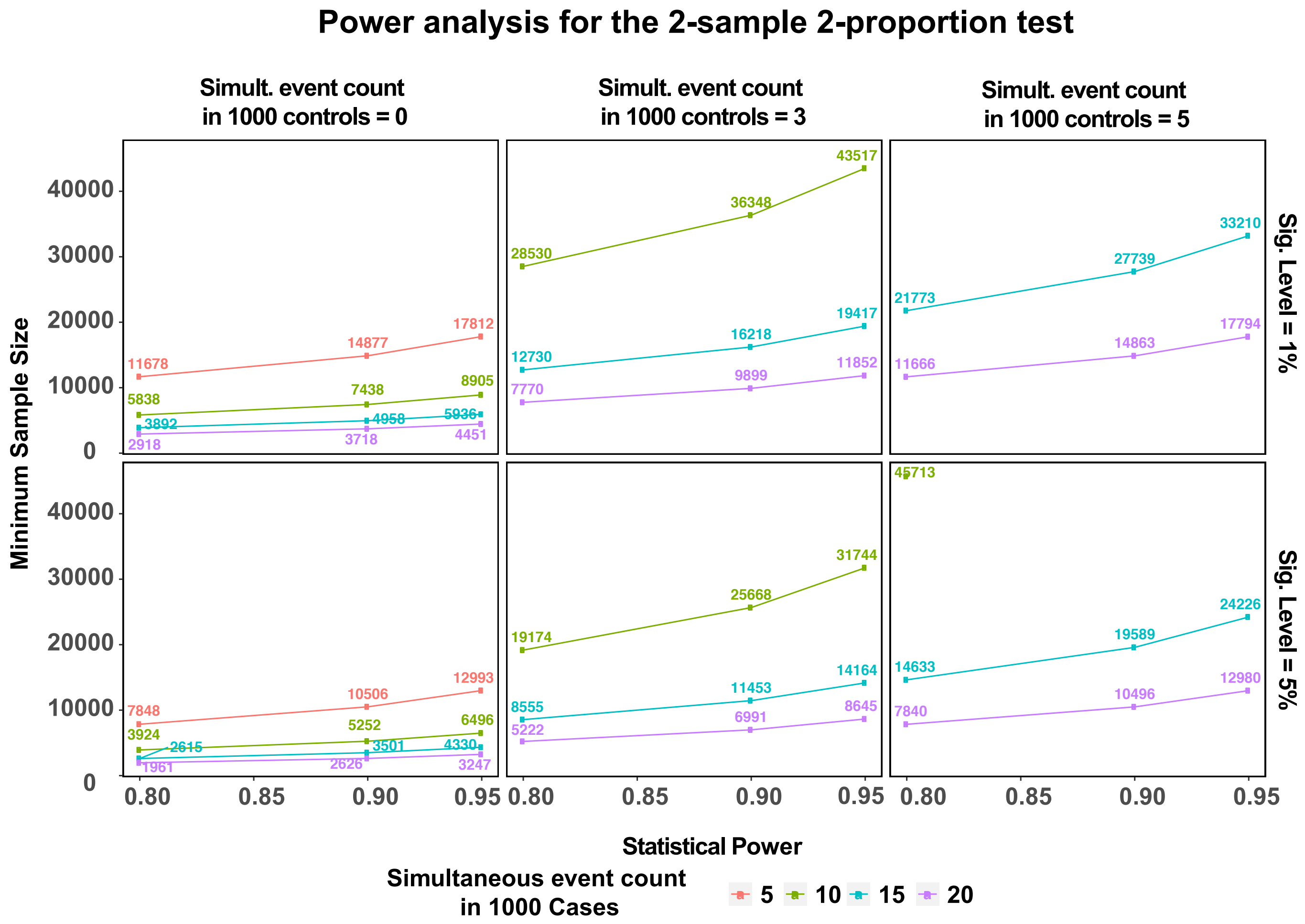

**Supp. Figure 14: Power analysis for 2-sample 2-proportion test to compare the frequencies of co-occurring events in cases and controls.** The panels along the X-axis show three specific frequencies of co-occurring events (0, 3, and 5) observed in 1,000 controls samples, while the panels along the Y-axis show the sample size requirements at 1% and 5% statistical significance thresholds. Values along the X-axis represent the statistical power achieved, and Y-axis denotes the sample size needed to achieve the corresponding power. Each line color represents four specific frequencies of simultaneous events (5, 10, 15 and 20 out of 1,000 samples) in cases. *For example*, to establish statistical difference between a co-occurring event that occurs 10/1,000 times in cases (green) and 3/1,000 times in controls (middle panel along the X-axis), it would take 19,174 samples to achieve a statistical power of 80% at 5% significance threshold (bottom panel along the Y-axis). The colors missing in some of the panels show that the sample size requirements are higher for such configurations to fit into this graph.

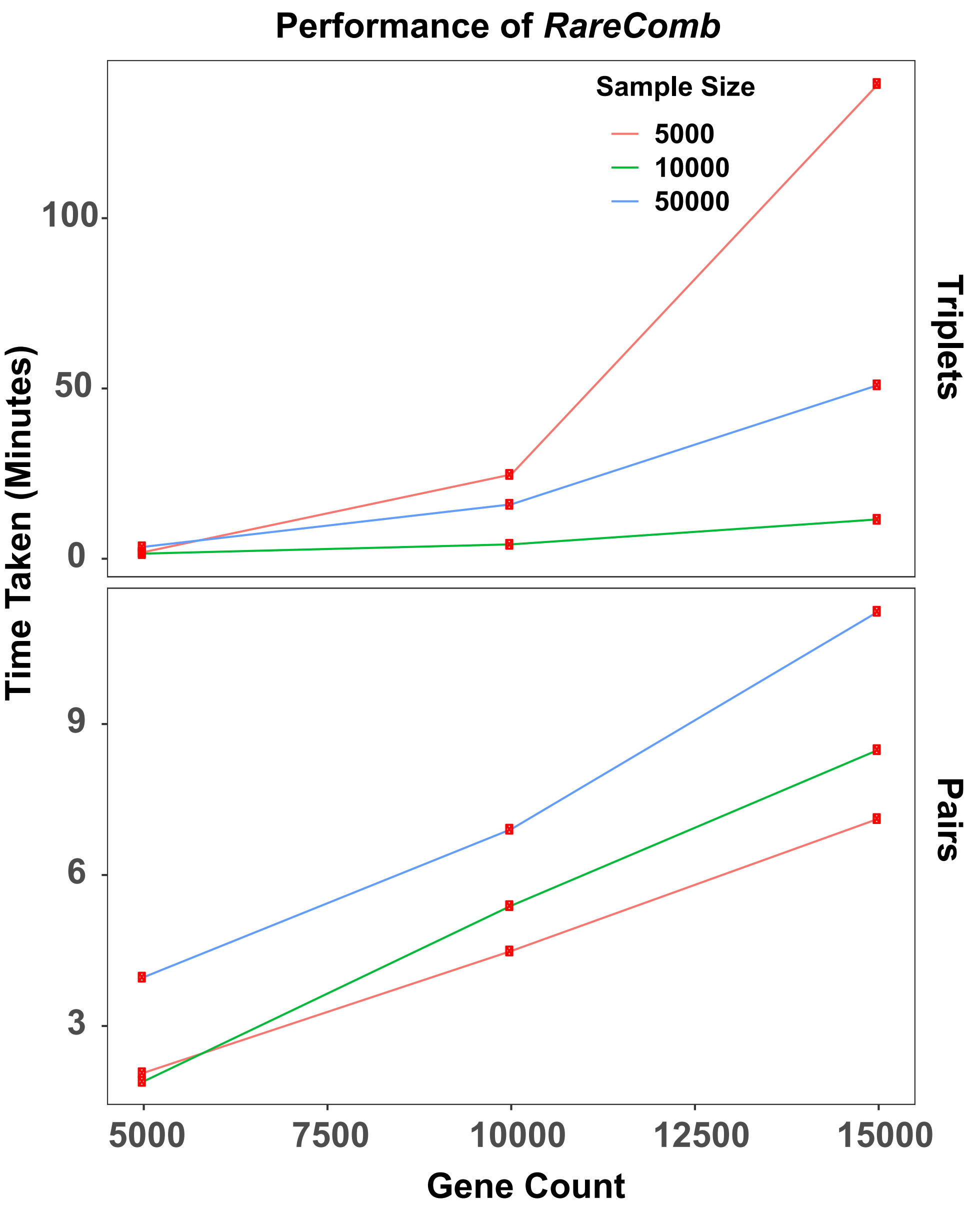

**Supp. Figure 15: Performance of *RareComb*.** Time taken by the pipeline to identify significant pairs and triplets using input files of various width and length is shown. The panels along the Y-axis represent the time taken to generate pairs versus triplets, and the Y-axis represent the time taken, in minutes, by *RareComb* to generate results. The values along the X-axis indicate the number of genes in the input file, and the line colors represent the sample size within the input files. As expected, an increase in the number of predictors is accompanied by the increase in the time taken by the method to generate pairs and triplets. Similarly, for pairs, the time taken increases with the increase in sample size. However, due to stochasticity in the input data and the complex relationship between the size of data under analysis and the minimum frequency threshold provided to the apriori algorithm, the method generated triplets faster with 50,000 samples compared to 10,000 samples. Notably, the method can generate results for pairs within 15 minutes and for triplets within three hours.
